# FlickerPrint: An Analysis Package for Measuring Interfacial Tension and Bending Rigidity of Biomolecular Condensates and Vesicles at Scale

**DOI:** 10.1101/2025.03.24.645013

**Authors:** Thomas A. Williamson, Jack. O. Law, Thomas Stevenson, Fynn Wolf, Carl M. Jones, Endre S. Tønnessen, Sushma N. Grellscheid, Halim Kusumaatmaja

**Affiliations:** Institute for Multiscale Thermofluids, School of Engineering, University of Edinburgh, Edinburgh, UK; Computational Biology Unit and Department of Biological Sciences, University of Bergen, Bergen, Norway; Department of Biosciences, University of Durham, Durham, UK

**Keywords:** biomolecular condensate, vesicle, flicker spectroscopy, interfacial tension, bending rigidity

## Abstract

**In Brief:** Williamson, Law *et. al*. present *FlickerPrint*, a computational analysis tool which can be used for measuring the interfacial tension and bending rigidity of soft fluctuating bodies, including biomolecular condensates, droplets or vesicles, from confocal microscopy images using flicker spectroscopy. This method is highly scalable so can be used to analyse the properties of whole populations of thousands of such soft bodies.

**Motivation:** Biomolecular condensates play fundamental roles in sub-cellular organisation and it is well-known that the composition of condensates can affect their function. Measuring the condensates’ mechanical properties (for example, interfacial tension and bending rigidity) can aid the understanding of their biomolecular composition and cellular functions. However, measuring the properties of individual condensates under physiological conditions is very challenging and cumbersome to scale to the population level using traditional methods. To overcome these issues, we have developed a software package to run flicker spectroscopy analysis of condensates at scale, to determine their interfacial tension and bending rigidity. At the same time, *FlickerPrint* can be harnessed to analyse other soft, fluctuating bodies such as lipid vesicles.

**Summary:** Accurate measurement of the mechanical properties of biomolecular condensates is an essential step in understanding their behaviour within cells. We present *FlickerPrint*, an open-source Python package to determine the interfacial tension and bending rigidity of thousands of biomolecular condensates using flicker spectroscopy by analysing their shape fluctuations in confocal microscopy images. We detail the workflow used by *FlickerPrint* to scale up these individual measurements to the population level and the computational requirements to run *FlickerPrint*. We provide examples of experiments in live cells and *in vitro* which are suitable for analysis with *FlickerPrint* as well as scenarios where the package cannot be used. Using these examples, we show that the results obtained from the analysis are robust to changes in the imaging setup, including frame rate. This implementation enables a step-change in the capability to measure two key properties of biomolecular condensates, the interfacial tension and bending rigidity. Moreover, the tools in *FlickerPrint* are also applicable for analysing other soft, fluctuating bodies, which we demonstrate here using vesicles.

## 5 Introduction

Biomolecular condensates are phase-separated, liquid like droplets which exist across a wide range of different environments, both *in-vitro* as synthetic droplets and in cells as cellular subcompartments such as stress granules, nucleoli and p-bodies, where they regulate important functions from stress response to RNA transcription^1–5^. One way to help to understand the behaviour of condensates is by measuring their mechanical properties, a set of generalised parameters which govern the condensate’s mechanical response, from shape changes to droplet coalescence, wetting onto membranes and production of biomolecules^2,6–8^. In turn, the mechanical properties of a condensate are governed by its internal biomolecular interactions^9,10^.

Much work has focused on the bulk properties of condensates, showing that they are network fluids and that viscoelasticity plays an important role in governing their dynamics^11,12^. However, the condensate interface has also been shown to play an important role in modulating chemical reactions, wetting and the liquid-to-solid ‘ageing’ transition which some condensates appear to undergo^13–18^. The interfacial properties of condensates depend on the interactions between the molecules in the dense and dilute phases. Therefore, while *in-vitro* techniques can provide highly accurate measurements of properties such as interfacial tension, these results may not translate directly to *in-cellulo* assays^19–21^. Non-invasive techniques allow for interfacial properties to be measured in live cells, pioneered with estimates of interfacial tension from the time taken for droplets to coalesce^22^. However, since this technique relies on the observation of relatively rare coalescence events, it can be difficult to conduct at scale. More recent work has estimated the mechanical properties of coacervates by fitting a statistical model to their size distribution^23^. While this can provide a useful population-level estimate of the average mechanical properties, it is not able to provide individual-level measurements to understand how parameters vary across an entire population.

To overcome these issues, we present *FlickerPrint*, a comprehensive, high-throughput Python package which implements the flicker spectroscopy method for measuring interfacial tension and bending rigidity (a measure of the elastic bending deformation of the interface) of condensates and other soft, fluctuating bodies such as vesicles^6^. *FlickerPrint* non-invasively measures the mechanical properties of condensates or vesicles *in vitro* or in live cells using confocal microscopy images. These measurements are repeated at scale to produce parameter distributions for whole populations of the objects of interest, with individual-object resolution. The ability to collect measurements on a large scale using widely available microscopy setups will enable these assays to be carried out more routinely, so that the behaviour of soft bodies such as condensates and vesicles can be better-understood.

The flicker spectroscopy method determines the interfacial tension *σ* and bending rigidity *κ* by measuring the shape fluctuations of biomolecular condensates or vesicles in confocal microscopy images^6,22,24^. By equating the Helfrich-type energy penalty of the shape fluctuations at the interface to the thermal energy of the object of interest, we can derive an analytic power spectrum of the amplitude *ν* of the *q*^th^ fluctuation mode

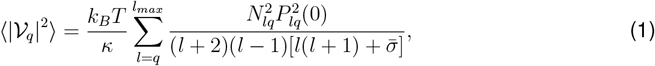

where *N*_*lq*_ *P_lq_* is the normalised Legendre Polynomials, 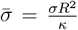 and *R* is the radius of the object^25,26^. Taking *l*_max_ = 75 is typically sufficient for the sum to converge. Fitting this power spectrum to the experimentally determined fluctuation amplitudes allows the interfacial tension *σ* and bending rigidity *κ* to be determined as free parameters^6^. The flicker spectroscopy technique was originally developed in the context of vesicles^27,28^ and red blood cells^29,30^, and we recently demonstrated that it can be used to study biomolecular condensates^6^.

In this work, we outline the workflow used by *FlickerPrint* and describe the types of systems which are suited to analysis with the package. We then detail the parameters which can be measured using *FlickerPrint*, including interfacial tension, bending rigidity and condensate shape, before describing the technical considerations when collecting microscopy videos for analysis. Finally, we detail how *FlickerPrint* can be installed and the computational requirements for running the package.

## 6 Results

### 6.1 Overview of the *FlickerPrint* package

*FlickerPrint* takes confocal microscopy images of condensates or other soft bodies (either in live cells or *in vitro*) and uses the flicker spectroscopy method which we have previously described to determine their interfacial tension and bending rigidity, amongst other properties^6^. The package is available on all major platforms (Windows, Linux, macOS) and can be run in either desktop or High Performance Computing (HPC) environments.

A flowchart showing the basic workflow of *FlickerPrint* is shown in figure 1A; the full control logic is shown in figure S1. The workflow can be split into three main stages: (i) location of objects of interest, (ii) extraction of the Fourier terms from the boundary fluctuations, and (iii) fitting of the theoretical spectrum to determine interfacial tension and bending rigidity.

**Figure 1.**
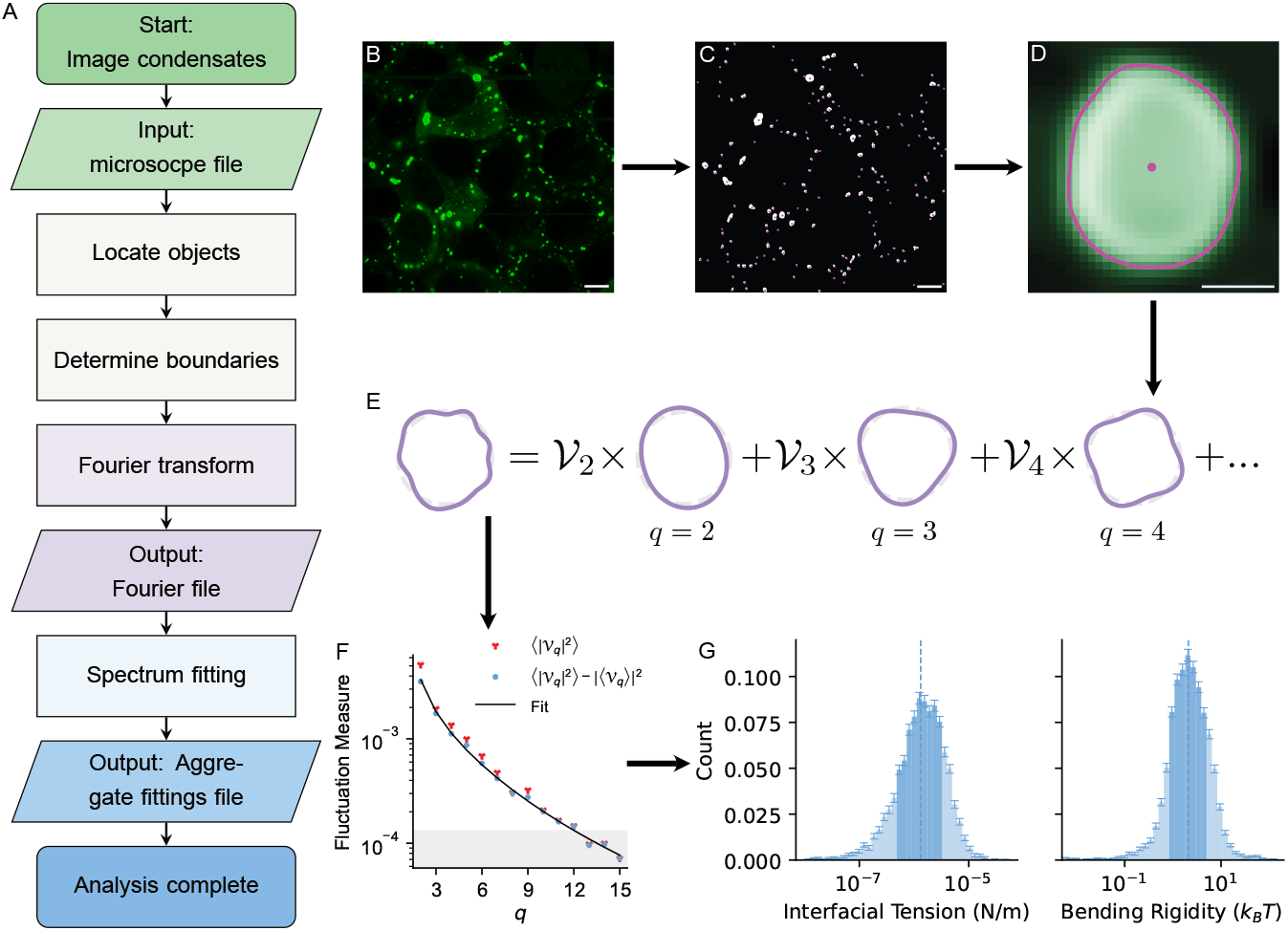
The workflow used by *FlickerPrint*. (**A**) A flowchart outlining the key stages of the analysis. Colours indicate the stage of analysis; condensate detection and location (green), extraction of fluctuation Fourier components (purple) and power spectrum fitting (blue). (**B**-**G**) Illustrations of the key stages. (**B**) A raw micrograph input into *FlickerPrint* showing stress granules induced in U2OS cells using sodium arsenite. G3BP1 is fluorescently labelled. (**C**) The centres of condensates are located (magenta dots) using the Difference of Gaussians method and their approximate shape is determined using a flood fill (white regions). (**D**) The boundary of each condensate is determined as the maximal intensity gradient from the centre of the condensate. (**E**) The magnitudes *ν*_*q*_ of the Fourier modes of the condensate boundary are determined for each condensate in each frame. (**F**) A power spectrum of the boundary fluctuations is produced and corrected for the base shape of the condensate. The interfacial tension and bending rigidity are the best fit parameters of equation 1 to the experimental spectrum. (**G**) The boundary determination, Fourier Transform and spectrum fitting steps are repeated for all condensates in the experiment to build up a population distribution of interfacial tension and bending rigidity. Scale bars: 10 *µ*m (B,C); 1*µ*m (D).

In the first stage, the objects are located and their boundary is determined. A Difference of Gaussians (DoG) algorithm is used to detect condensates or vesicles as high intensity regions relative to their local environment (Figure 1B)^31^. This allows for substantial variations in background intensity between cells to be accounted for, when applied in an *in cellulo* context. The size of the objects of interest to be found can be tuned by configuring the width of the Gaussian filters used, which can be useful to ensure analysis of other cellular structures is avoided. The objects are tracked through frames in the video using a custom tracking algorithm (SM1), so that their shape fluctuations can be measured with time. Once the objects of interest have been located, a flood fill is used to determine their approximate extent and to draw a bounding box around them. Next, the boundary of the object must be determined. In the microscopy frames, condensates appear to have a diffuse boundary. As such, the condensate boundary is taken as the maximal intensity gradient in the radial direction from the centre of the condensate (SM2). For vesicles, the local maximum intensity of the image is used instead. This results in a function of the radius of the object at a polar angle *φ* about its centre point (Figure 1D). Typically, the boundary can be determined to within 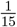 of a pixel (SM3).

In the second stage, the boundary is normalised by the average radius of the object and broken into its constituent modes by taking a Fourier Transform (Figure 1E) to yield the amplitude *ν* of each Fourier mode *q*. Typically, modes 2 ≤ *q* ≤ 15 are used (*q* = 0 represents changing the size of the object and *q* = 1 represents translational movement, both of which are accounted for in earlier stages of the analysis). For modes higher than *q* = 15, the amplitude of the mode is often close to the resolution limit of the boundary detection (Figure 1F), though the number of modes measured can be adjusted, if applicable.

Stages 1 and 2 are repeated for all of the input microscope videos. Intermediate files are produced which contain the locations, sizes and boundary Fourier components of all objects in each frame of the videos.

The third main stage of the *FlickerPrint* workflow is fitting to the power spectrum. Simple Newtonian liquids have a spherical time-averaged base shape so taking a Fourier transform of their boundary should only yield their thermal fluctuations. However, condensates and vesicles do not always have a spherical base shape. The shape of a condensate in a given frame is comprised of two components: the time-averaged ‘base shape’ of the condensate ⟨*ν*_*q*_⟩ and the instantaneous thermal fluctuations |*F*_*q*_| on top of this shape.

Therefore, the contribution of ⟨*ν*_*q*_⟩ is removed according to^32^

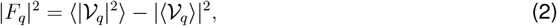

to leave only the fluctuating contributions |*F*_*q*_ |^2^. Equation 1 is fitted to the corrected experimental spectrum using a least-squares fit (figure 1F) with a custom minimisation function (SM4) to ensure that all orders are fitted with equal weighting. Interfacial tension *σ* and bending rigidity *κ* are the only free parameters in the fit.

A final ‘aggregate fittings’ output file is produced, containing the interfacial tension and bending rigidity of all objects in the experiment, together with additional useful properties such as their mean radius and mean intensity. Population distributions such as those in Figure 1G can be produced using *FlickerPrint* ‘s built-in graphing tools or any other standard statistical analysis toolkit.

### 6.2 Experiments suited to analysis with *FlickerPrint*

*FlickerPrint* takes microscopy videos as input, showing bright condensates or vesicles fluctuating against a dark background. Therefore, while our initial application of the method was to stress granules in live U2OS cells, FlickerPrint can be used to characterise a variety of *in-cellulo* and *in-vitro* systems of both condensates and vesicles^6,24^. Figure 2 shows four systems: a system of stress granules induced in U2OS cells using sodium arsenite (A), an *in vitro* system of NPM1 droplets using a dextran crowding agent (B), a system of solid fluorescent polystyrene particles (C) and a system of coacervate-core vesicles, formed of peptides with a 2,2’-thiobis(ethylamine) spacer and two tyrosine-phenylalanine stickers (YFsFY) (D)^24^. The condensates and polystyrene particles appear as bright, filled regions (Figure 2 E-G) whereas vesicles appear as high-intensity outlines (Figure 2 H); both intensity profiles can be used to determine the fluctuation spectrum of the objects of interest.

**Figure 2.**
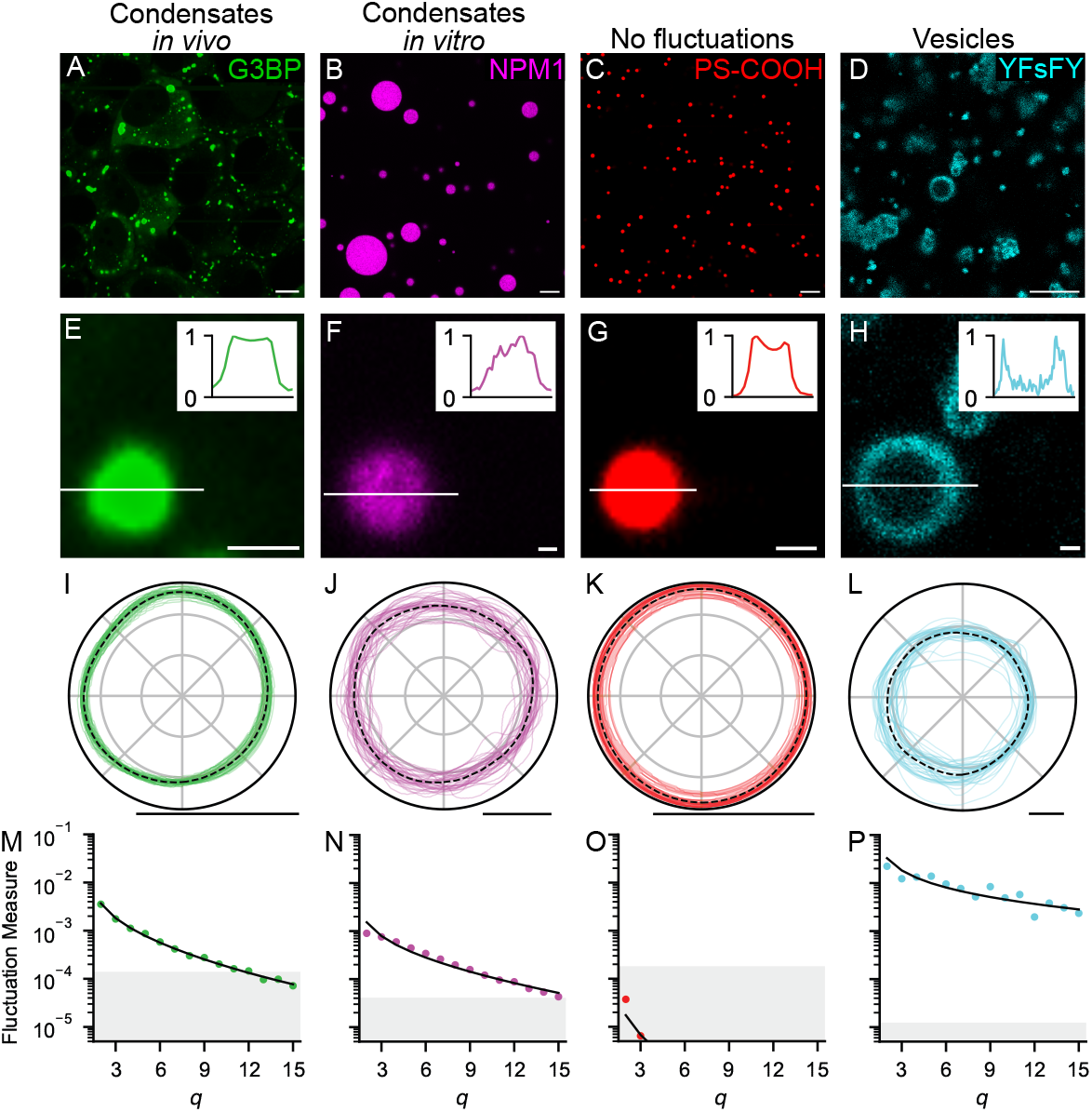
Examples of the suitability of different experimental setups for analysis with *FlickerPrint*. (**A**-**D**) confocal microscopy images of four different systems; stress granules in live U2OS cells (A), synthetic NPM1 condensates with a Dextran crowding agent (B), fluorescent carboxyl-functionalised polystyrene particles (C) and vesicles (D). The fluorescently labelled species is given in each sub-figure. (**E**-**H**) Zoom in of a single object of interest from (A-D). Insets give the intensity profile along the line shown. (**I**-**L**) Boundary fluctuations of the object shown in (E-H) with time. The black dotted line shows the time-averaged base shape of the condensate or vesicle. For clarity, only the first 40 fluctuations are shown. (**M**-**P**) The shape corrected power spectrum for the fluctuations (dots) of the objects in (E-H) with the fit from the theoretical power spectrum shown as a black line. Fluctuation measure is normalised to the object’s mean radius. The shaded grey region denotes where the total fluctuation amplitude is too small to be resolved reliably (typically *<* 0.067 pixels). **D**,**H**,**L**,**P** are reproduced with permission using data from^24^. Scale bars: 10 *µ*m (A-D); 1*µ*m (E-L).

The fluctuations of each object are shown in figure 2 I-L. Where the time-averaged base-shape is non-spherical (Figures 2 I, L), this contribution can be decoupled from the fluctuations using equation 2 ^32^. The fluctuations of stress granules, NPM1 condensates and vesicles (Figures 2 I, J, L) are all large, compared with their radius. As such, the fluctuation spectra are almost entirely above the resolution guide (Figures 2 M, N, P). Therefore, these species are suitable for analysis with *FlickerPrint*. In contrast, the ‘fluctuations’ seen in Figure 2 K are caused by movement of the bead into and out of the imaging plane, causing an apparent change in size but no change in shape. This does not impact the power spectrum. The lack of visible shape fluctuations means that the entire fluctuation spectrum of the polystyrene beads (Figure 2 O) is below the resolution guide and so interfacial tension and bending rigidity cannot be determined using *FlickerPrint*, as expected for solid particles.

In addition to the objects having visible thermal fluctuations, they must also exist in thermal equilibrium and in ‘free space’. This means that condensates which are undergoing fusion, or are wetted onto other cellular apparatus (or their container in an *in-vitro* setting) cannot be analysed. The boundary of the condensate must also be able to be described as a continuous radial function *D*(*φ*) of the polar angle about its centre-point, as shown in 1 D. Figures 3 A, B show examples of stress granules in live cells which have been rejected because their boundary is not continuous. Often, where condensates are not in equilibrium or have external forces acting upon them, their shapes cannot be described as a continuous radial function (Figure 3 A shows two stress granules fusing, for example). Condensates which have undergone ageing to form solid-like aggregates also often fail this shape filter, however since they are solid-like, they are unlikely to show thermal fluctuations above the resolution threshold of the microscope anyway.

**Figure 3.**
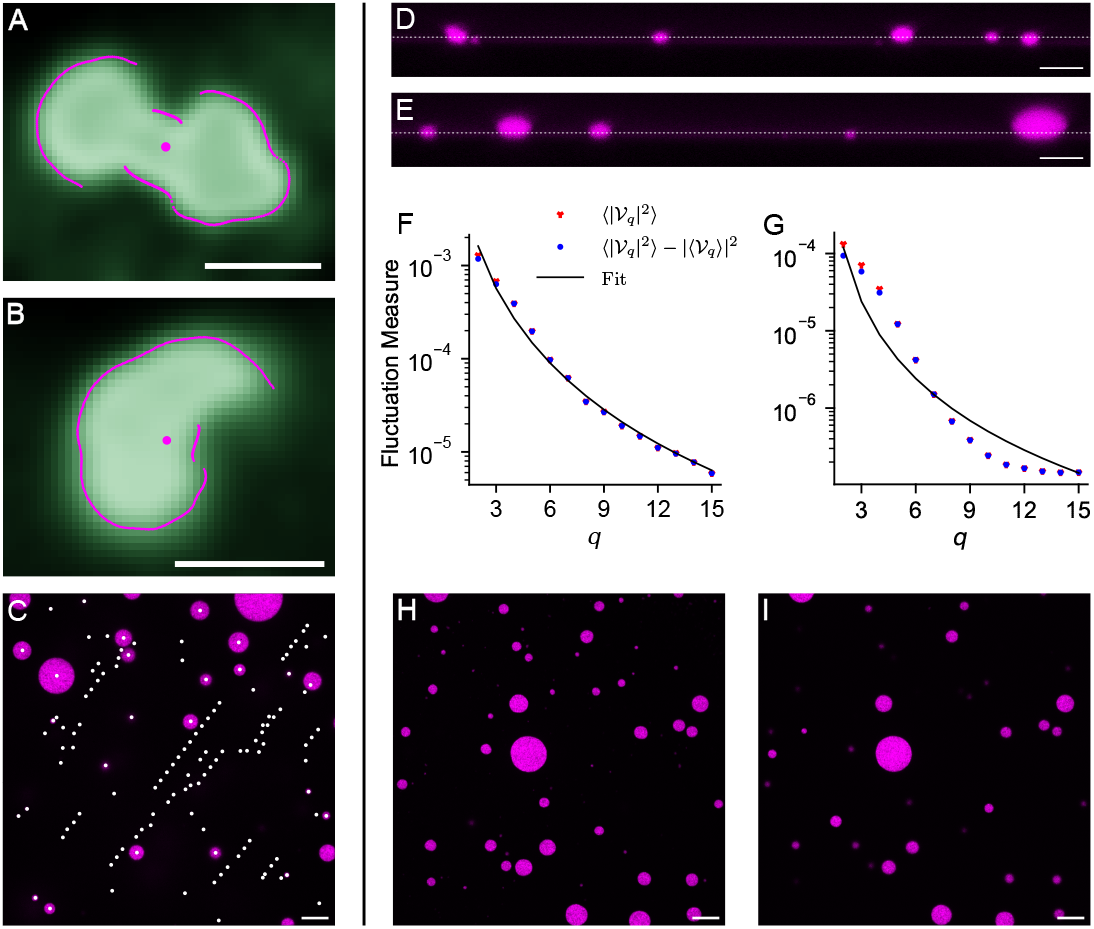
Ensuring that images are captured appropriately is essential for successful analysis using *FlickerPrint*. (**A**,**B**) Examples of stress granules which have not passed the boundary filter as their boundary cannot be expressed as a continuous radial function (magenta line) about the centre. (**C**) Image showing the effect on condensate tracking in FlickerPrint when condensates move too quickly. The effect is simulated by moving the microscope field of view in between frames. White dots show the location of ‘new’ condensates found across all frames in the video. (**D**) Side profile (x,z) confocal microscopy image of an *in-vitro* system of synthetic NPM1 (magenta) condensates with a dextran crowding agent, showing condensates of approximately the same size settled onto a slide. The white dashed line shows an example imaging plane passing close to the equator of the condensates. (**E**) Similar to (D) but for condensates which are not of an even size. In this case, it is not possible for the imaging plane to pass within ± 0.4 radii of the equator of every condensate. (**F**) Example spectrum for an NPM1 condensate which has been imaged close to the equator. (**G**) Similar to (F) but for a condensate which has been imaged far from the equator, leading to a very high fitting error. (**H**) Top down (x,y) confocal microscopy image of the *in-vitro* NPM1 system, where the imaging plane has been selected to maximise the number of condensates imaged. (**I**) Similar to (H) but where the imaging plane has been moved up to capture the equatorial region of the larger condensates. Micrographs in positions similar to both (H) and (I) are required to produce suitable fluctuation spectra for both large and small condensates. Scale bars: 1*µ*m (A,B); 10*µ*m (C-E,H,I).

The final requirement of experiments to be analysed using FlickerPrint is that the spatial position of the objects of interest should be relatively stable. *FlickerPrint* tracks the location of objects to account for small lateral movements in the XY imaging plane, up to a displacement of 15 pixels per frame (SM2). If the objects move by more than 15 pixels per frame (as demonstrated in Figure 3 C, where movement was simulated by moving the microscope field of view between frames), the objects may be lost by the tracking algorithm. If they are re-found far from their original position in subsequent frames, they may be tracked as a new object, which may lead to the same object being analysed multiple times, impacting the output parameter distributions.

It is also important to ensure that the z position of condensates remains stable, relative the confocal microscopy imaging plane. In particular, equation 1 assumes that condensates have been imaged at their equator, though the uncertainty in a condensate’s interfacial tension and bending rigidity is *<* 20% when the imaging plane is within 0.34 radii of the condensate equator^6^. Typically, this condition is met for condensates in live cells on a single microscope plate. Special care must also be taken when measuring the properties of *in-vitro* condensates. It is preferable to allow *in vitro* condensates to settle onto the bottom of their container, to stabilise their Z position, though the surface should be treated so that the condensates do not wet onto the surface. If the objects of interest are of approximately uniform size, it is possible for the imaging plane to pass within 0.34 radii of the equator of all condensates (Figure 3 D). However, when *in-vitro* condensates are settled onto a surface but are not of approximately uniform size, choosing an imaging plane which maximises the number of condensates in the frame (Figure 3 E) will mean that larger condensates are imaged more than 0.34 radii from their equator. When condensates are imaged away from their equator, the shape of their fluctuation spectrum changes substantially (Figures 3 F, G), resulting in a significantly higher fitting error 0.073 vs. 1.093) and leading to a poor estimate of the condensate’s mechanical properties. To combat this issue, images can be taken in two different planes (Figures 3 H, I) which are suitable for imaging the equators of smaller condensates and larger condensates respectively. At the analysis stage, the condensates can be filtered according to size, to prevent double-counting.

### 6.3 Parameters which can be measured

For experiments which are suitable, *FlickerPrint* can be leveraged to measure multiple properties of a population of condensates or vesicles. These properties are measured at the individual level and repeated at scale to provide parameter distributions at the population level. The analysis is parallelised across multiple microscope videos, where appropriate. The parameters that can be measured are summarised in Figure 4 and are broadly grouped into three main categories: properties which are determined by the time-averaged state of the object of interest, those which are determined by the fluctuation spectra of the object and additional properties which can be used to filter the population dataset.

**Figure 4.**
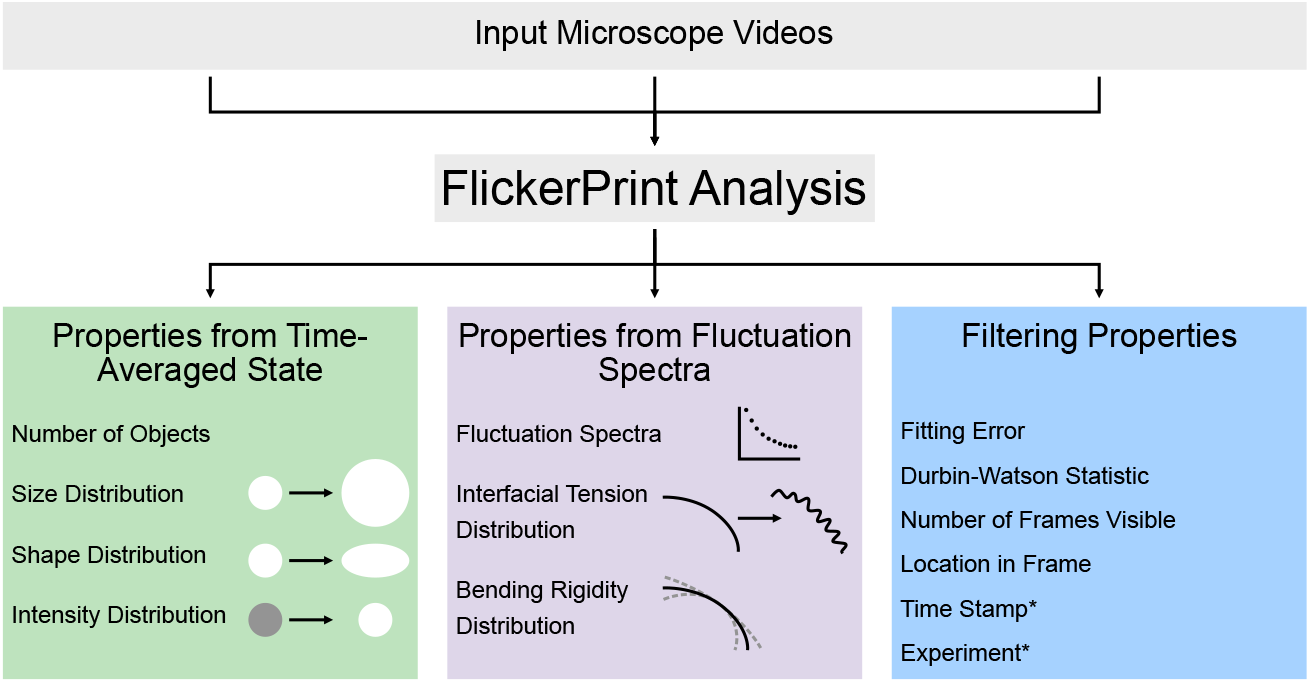
*FlickerPrint* allows the mechanical properties of condensates and vesicles to be determined at scale. The properties can be grouped into three broad categories: those which are determined from the time-averaged state of the objects of interest, those which are determined from their thermal fluctuations and additional properties which can be used for filtering population datasets. *Time stamp and experiment name are supplied from external metadata.

The first category of properties are determined from the time-averaged base shape of the condensate or vesicle and include basic measurements such as the number of condensates measured and their size and intensity distributions. These are regularly quantified in the literature and can be measured by existing packages^33^. Together, they can provide valuable information about the propensity of a system to phase separate^34,35^. FlickerPrint allows this analysis to be easily parallelised across multiple microscope videos from the same assay. However, *FlickerPrint’s* capabilities go beyond these standard measurements; less readily characterised is the shape distribution of populations of condensates and vesicles. Condensates are understood to take on irregular shapes when they age to form solid-like aggregates or when they are deformed by other cellular components such as fibrils^14,36,37^. However, there is evidence that stress granules can also take on non-spherical base shapes, which may be evidence of their viscoelastic behaviour^6^. In the context of vesicles, the base shape may provide information on the membrane area to volume ratios of the system^38^.

The second set of properties are those determined from the fluctuation spectrum of the objects of interest. These include the raw fluctuation spectra themselves, as well as the interfacial tension and bending rigidity derived from the spectra. Flicker spectroscopy has previously been used to successfully measure the properties of individual vesicles and red blood cells^27–30,39^. However, to the authors’ knowledge, *FlickerPrint* is the only software package which can measure interfacial tension and bending rigidity of individual biomolecular condensates and vesicles at scale to allow for population-level analysis. Therefore, both the mean properties and shape of the property distributions can be determined and compared, to understand how properties interact to influence condensate behaviour. Recent estimates of mean interfacial tension from the size distribution of CPEB4Δ4_NTD_ coacervates give values of 1 − 1.5*µ*N*/*m, in broad agreement with the values which we have previously found for stress granules in U2OS cells^6,23^.

The final set of properties are those which are produced as a result of the data collection and analysis, and can be used for filtering when visualising population-level parameter distributions. These include the fitting error and Durbin-Watson statistic (a measure of the autocorrelation between residuals associated with fitting equation 1 to the fluctuation spectrum). Examples of spectra with good and poor fitting error and Durbin-Watson statistic are shown in Figure S2. The number of frames which an object of interest is visible for can also be used as a filter to ensure that only objects with enough shape fluctuations to be properly time-averaged are included in the parameter distributions. In addition, the image timestamp, a user-defined experiment name (both supplied externally) and the location of objects within the frame can be used to aid in the comparison of parameters measured in different assays. A full list of parameters which can be measured is available in the *FlickerPrint* documentation^40^.

### 6.4 Imaging Considerations

*FlickerPrint* is compliant with the Euro-BioImaging FAIR standard and can accept image files in one of the following formats: .*ims*, .*lif*, .*ome*.*tif[f]*, or .*tif[f]* ^41^. It may be possible to use other Bioformats compliant file-types, but these are untested^42^. It is recommended that unsupported file types are converted to .*tif* or .*ome*.*tif* files prior to analysis. In addition, some of the above file-types may be missing important metadata, and must be handled carefully; .*tif* files do not contain the pixel-size in their metadata, so this must be recorded manually from the microscope and provided explicitly to *FlickerPrint via* the analysis configuration file.

The measured interfacial tension and bending rigidity of a single population of condensates can span multiple orders of magnitude^6^. Therefore, a large sample size is required to sufficiently map out the parameter distribution at the population level. For a single assay, it is generally best to generate at least 10 videos by moving the microscope field of view around the sample, focusing on areas with a high density of condensates. A final count of above 1000 analysed condensates may require up to 50 input videos, depending on the condensate density and the fraction of those condensates that pass the filtering steps.

In addition to the total number of objects of interest that are analysed, it is also important to try to maximise the total number of frames each object is visible for; each object should ideally be visible for at least 200 frames, with ∼1000 frames being optimal. This ensures that the magnitudes of the fluctuations can be properly averaged, as required by equation 1. The frame rate used to capture the micrographs should be high enough to ensure that the imaging timescale is much smaller than the timescale of condensate ageing or any other changes in the mechanical properties. However, provided this condition is met, the frame rate used does not substantially affect the measured properties of the condensates, as demonstrated in Figures 5 A, B and Table S1; the mean interfacial tension and bending rigidity deviate by 0.043N*/*m and 0.03*k*_*B*_*T* (corresponding to *<* 0.3 and *<* 0.06 geometric standard deviations of the population distributions) respectively, when comparing condensates which are found and pass the necessary filters in all four analyses. When stress granules which were not found across all four analyses are included, the mean interfacial tension and bending rigidity deviate by 0.12N*/*m and 0.19*k*_*B*_*T* (corresponding to *<* 0.5 and *<* 0.3 geometric standard deviations of the population distributions) respectively (Table S2), indicating that the differences can largely be attributed to an increased number of condensates being found and successfully tracked at higher frame rates.

**Figure 5.**
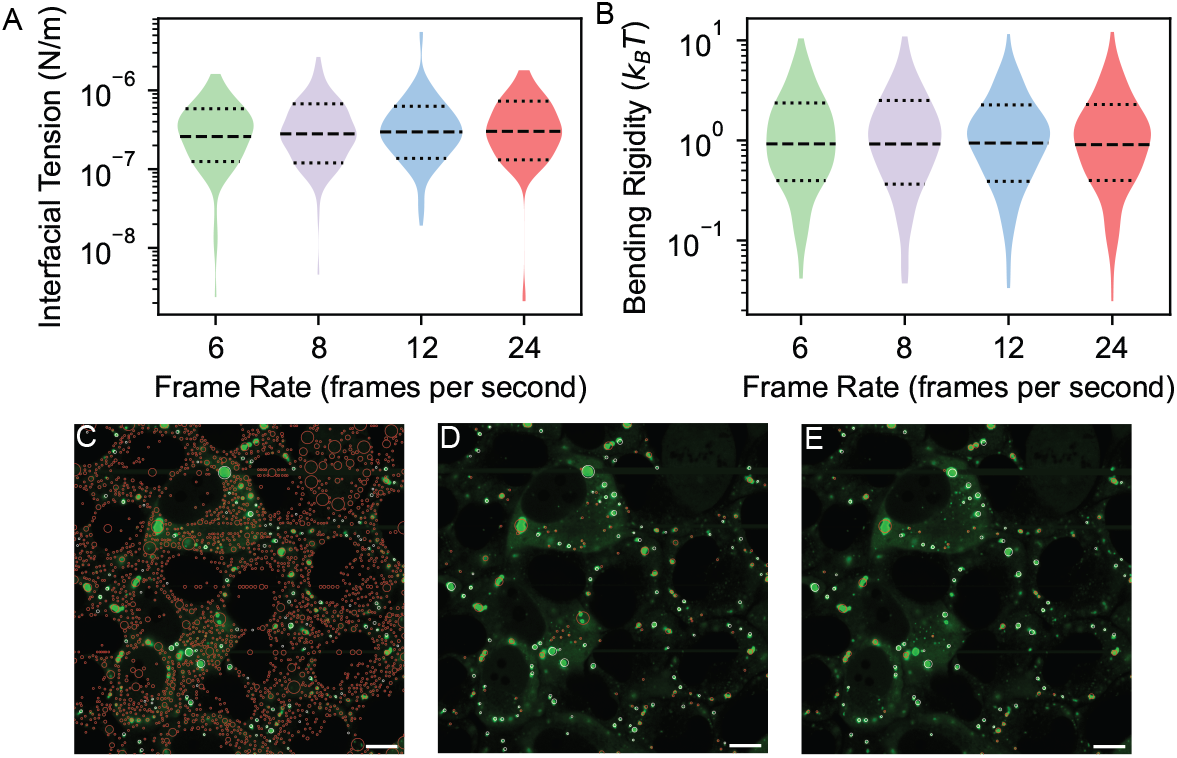
Ensuring that microscope videos are captured and imaging parameters are configured effectively leads to the highest chance of successful use of *FlickerPrint*. (**A**,**B**) Violin plots showing the distribution of interfacial tension and bending rigidity for stress granules induced using sodium arsenite in U2OS cells, analysed at effective frame rates of 6, 8, 12 and 24 frames per second. Two videos were taken, each at 24 frames per second, and every *n*^th^ frame was analysed in order to produce the effective frame rates shown. (**C**-**E**) The effect of imaging parameters on the number of found condensates; microscope images of stress granules induced in U2OS cells, with the location of condensates found by *FlickerPrint*. Red circles show condensates which go on to be rejected by the boundary detection algorithm in this frame; white outlines show condensates which are accepted. The images show the effect of changing the minimum intensity parameter; (C-E) show intensity thresholds which are too low, optimal and too high respectively. Scale bars: 10*µ*m (C-E).

To account for differences in contrast, size and intensity of objects of interest in the videos, the directory of each experiment contains a configuration file which allows key imaging parameters to be adjusted. In particular, four parameters control the approximate maximum and minimum size of the detected objects, their minimum intensity and the flood fill used to determine their approximate extent. Since these parameters are used to locate objects of interest in each frame and do not directly impact the determination of the object boundaries, the analysis is relatively robust to their variation. When parameters are within their optimal range (the approximate range of values which maximise the number of objects found whist minimising the number which do not go on to pass the boundary continuity requirement), interfacial tension and bending rigidity vary by *<* 0.42 and *<* 0.52 geometric standard deviations respectively (Figure S4). A full discussion of the effect of imaging parameters on the returned property distributions is provided in SM5.

Once objects of interest have been located, their boundary is determined independently. Boundary detection is controlled by a single configurable parameter which acts to smooth the input images account for microscope noise (SM5). Typically, this parameter should be kept as close to 1.0 as possible, whilst minimising the number of objects where the detected boundary is not continuous.

Nevertheless, it is important that optimal imaging parameters are selected to maximise the number of condensates which can be analysed, whilst minimising the number of false detections which increase computational expense (Figure 5 C-E). To assist with selecting the optimal parameters for a given set of images, *FlickerPrint* contains a Bayesian optimisation tool. This tool implements a lexicographic optimisation to maximise the number of condensates found whilst minimising the number which do not go on to pass the later boundary filters (SM6)^43,44^.

### 6.5 Running *FlickerPrint*

*FlickerPrint* is available as a Python package for all major operating systems (macOS, Windows, Linux) and can be installed directly through PiP; the package is open-source and is also available on GitHub^45^.*FlickerPrint* requires Python 3.9-3.11 and Java 11.0.23 or later. Installation instructions and further guidance can be found in the documentation^40^.

*FlickerPrint* analysis can be scaled with core count, up to a maximum of one core per microscope image; the main analysis is executed through the command line using a single command. Data is analysed as an ‘experiment’, comprised of one or more microscope videos of the same biological system and defined by a single configuration file which contains the settings required for conducting the analysis. The experiment directory is also the location where intermediate and final output files are written to. All output files are saved in the HDF5 format, with data saved as standard c-type values, allowing them to be read by any standard statistical analysis software^46^. However, an additional Graphical User Interface tool is also provided (Figure 6). This allows for the production of 1D and 2D histograms so that the parameter distributions can be visualised and potential correlations between parameters can be found (see Figure 4 for examples of properties which can be measured using *FlickerPrint*). In addition, distributions from multiple experiments can be analysed simultaneously using violin plots, allowing for the comparison of the effect of different treatments or biomolecular compositions on the mechanical properties of the object.

**Figure 6.**
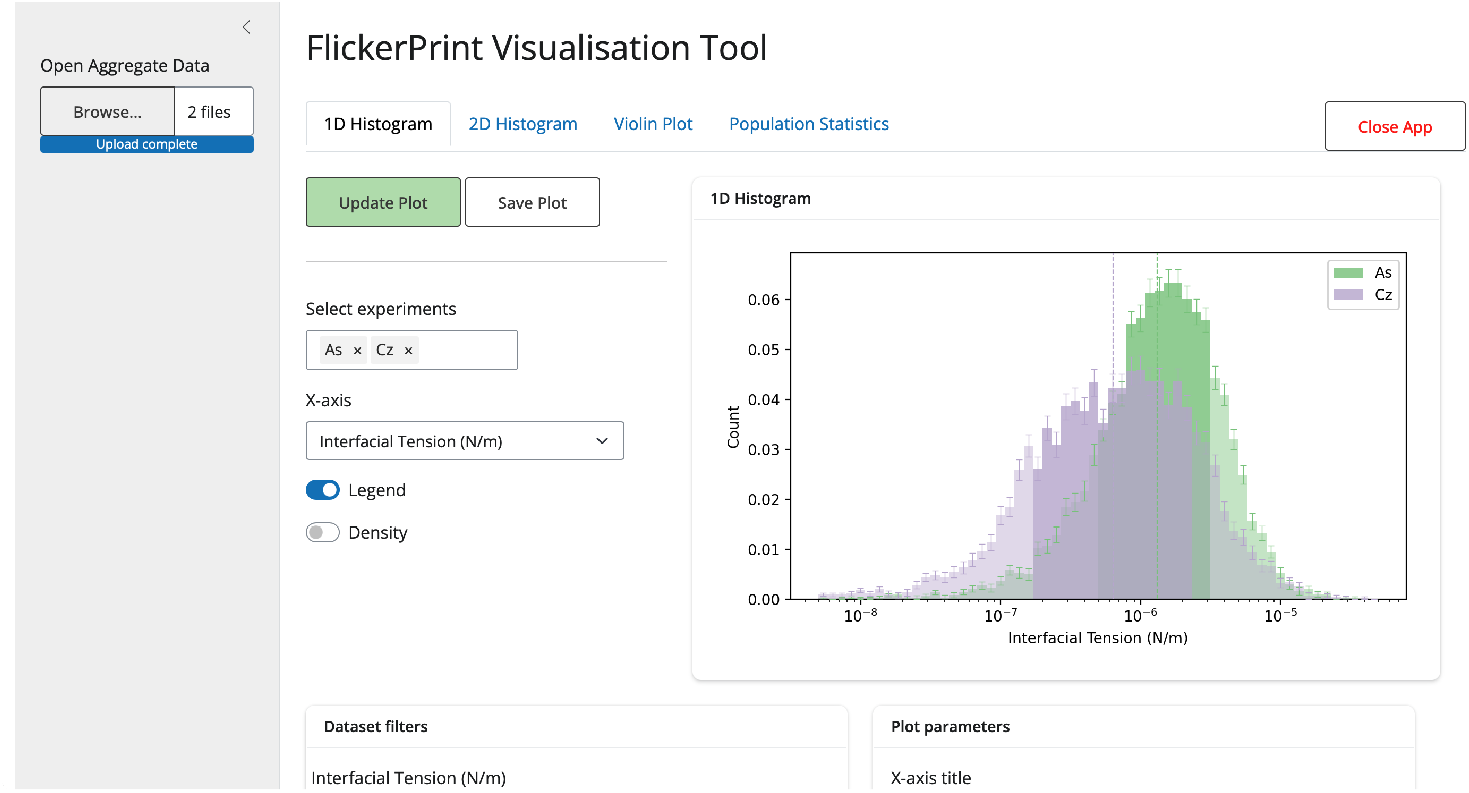
Screenshot of the Graphical User Interface application for producing plots of the parameter distributions from data output by *FlickerPrint*. Data used in these plots are reproduced with permission from^6^.

## 7 Discussion

In this work, we have presented *FlickerPrint*, an open-source Python package for undertaking flicker spectroscopy analysis of soft bodies at scale. We have demonstrated that this package can be used to measure the properties of biomolecular condensates and vesicles, both in live cells and *in vitro*. Principally, *FlickerPrint* is intended to measure interfacial tension and bending rigidity from shape fluctuations of the bodies. However, their time-averaged state can also be used to extract information on the size, number and shape of objects of interest. All of these properties can be measured as distributions at the population level, with individual-object resolution. We have also noted the requirements of experimental setups and microscope videos in order for them to be successfully used with *FlickerPrint*.

*FlickerPrint*, as a package for conducting flicker spectroscopy analysis, aims to make largescale surveys of the mechanical properties of condensates and vesicles more accessible. The scalable architecture used by *FlickerPrint* allows the package to be used in both desktop computing environments for testing and small-scale experiments, as well as High Performance Computing (HPC) environments for large-scale assays. The auxiliary tools for selecting optimal imaging parameters and analysing population-level data included in the package further improve its ease of use.

The confocal microscopy setup required for image collection for *FlickerPrint* is typically much more readily available in experimental biology laboratories than the setups required for du Noü y ring and similar experiments^19,20^. We anticipate that where changes are required to produce images of suitable quality for *FlickerPrint* (for example, by adjusting the imaging plane position), these should be easy to implement and should not require much more work than for collecting images for other purposes.

Measuring the coalescence time of condensates is another popular, non-invasive technique for estimating their interfacial tension^22,47^. While coalescence events themselves are typically quite fast, they are also relatively rare, meaning that these measurements provide a useful estimate of interfacial tension on long timescales for systems which do not evolve quickly in time. In contrast, the time scales required for image collection in *FlickerPrint* are of the order 1 minute, which may allow the properties of a population of condensates to be tracked with time. Dynamic Light Scattering experiments can also be used to measure the size distribution of condensates, from which an estimate of interfacial tension and bending rigidity at the population level can be determined by fitting a statistical model which accounts for differing protein confirmations at the interface and in the bulk of the condensate^23^. While this method can be used at scale, it can only provide an estimate of the mechanical properties of the system at the population level.

We believe that flicker spectroscopy acts as a complementary method to those described above for measuring the interfacial properties of individual biomolecular condensates, which can be scaled up to provide information about the mean and shape of the property distributions at the population level. *FlickerPrint*, as a package for completing the flicker spectroscopy analysis, makes the technique more accessible by implementing the analysis pipeline and handling the scaling required to measure the properties of populations of biomolecular condensates or vesicles.

### 7.1 Limitations of the Study

Throughout this work, we have outlined scenarios where it is not possible to use *FlickerPrint* (or the flicker spectroscopy method more generally) to determine the properties of a population of condensates. The first principle limitation is that condensates must not have solid morphologies, to ensure that their shape fluctuations can be measured (figure 2 O). We have previously shown that as stress granules age, fewer granules are able to pass the necessary filtering steps to complete the flicker spectroscopy analysis^6^. The second principle limitation is that condensates must exist in free space (so cannot be wetted onto containment vessels or cellular structures, for example) as these would provide additional energy contributions which would need to be accounted for in the fluctuation spectrum. However, even where it is not possible to accurately determine the interfacial tension and bending rigidity, *FlickerPrint* can still provide basic information on the size, circularity and fluorescence intensity distributions of the species being measured.

## 8 Materials & Methods

### 8.1 Condensates in Live Cells

U2OS ΔΔG3BP1/2 stably expressing GFP-G3BP1 were a gift from N Kedersha^48^. U2OS cells were cultivated in Dulbecco’s modified Eagle’s medium (DMEM) (Sigma-Aldrich, no. D5671) containing 10% fetal bovine serum, 100 µg/ml penicillin/streptomycin at 37 °C, 5% CO2. For imaging, cells were passaged and cultivated in 35 mm glass-bottom dishes (Ibidi, cat. no. 81158) or 18-well glass-bottom slides (Ibidi, cat. no. 81817) and stress granule condensates were induced via incubation in 200 uM sodium arsenite.

### 8.2 *In vitro* NPM1 Condensates and Polystyrene Particles

Reconstituted NPM1 condensates were formed in a buffer consisting of 20 mM Tris-HCl [pH 7.2], 250 mM potassium glutamate supplemented with 5 % Dextran. The final concentration of NPM1 protein in each experiment was 30 uM. Purified NPM1 and the NPM1 bacterial expression construct were provided by E Sprujit and was purified, labelled, and prepared as previously described^49^. Fluorescent (FITC) carboxyl-functionalised polystyrene particles were provided by E Sprujit and diluted in MQ water for imaging. Samples for FlickerPrint were imaged in functionalised 18-well glass-bottom imaging slides (Ibidi, cat. no. 81817), which were cleaned using a plasma cleaner and incubated overnight with 0.1 mg/mL PLL(2)-g[3.5]-PEG(2) (SuSoS, Dübendorf, Switzerland) dissolved in 10 mM HEPES (pH 8.0).

### 8.3 Microscopy

*FlickerPrint* -quality images for stress granule condensates in live cells, *in vitro* NPM1 condensates, and polystyrene particles were acquired using an Andor Dragonfly 505 spinning disk confocal system with a 100× 1.49 numerical aperture (NA) CFI SR HPApo total oil immersion objective using an iXon 888 or Zyla 4.2 PLUS sCMOS camera. XZ confocal micrographs of NPM1 condensates were acquired using a Leica TCS SP8 confocal microscope with a 100x 1.4 NA HC PL APO STED WHITE oil immersion objective. Distortion in the XZ micrographs was manually corrected via comparison to polystyrene particles of known size.

### 8.4 Population Filtering

In addition to the requirements placed on objects of interest (that objects can be located using the provided imaging parameters and that their boundary can be described by a continuous radial function) which act on a frame-by-frame and object-by-object basis, filtering can be applied at the population level. In this work, the following filters were applied to all population-level analyses: interfacial tension *σ* > 10^*−*10^N*/*m, continuous boundary in > 60% of frames, fitting error *ε* < 0.5 and |*ε* [*F* (*σ, κ*)] − *ε* [*F* (*σ*)] | > 0.03, where *ε* [*F* (*σ, κ*)] − *ε* [*F* (*σ*)] is the difference in fitting error between models which include contributions from both interfacial tension and bending rigidity or interfacial tension only.

## 9 Acknowledgements

We thank E. Spruijt, Radboud University for reagents and helpful discussions, N. Kedersha, Harvard University (retired) for reagents, N.A. Yewdall, University of Canterbury for support regarding the *in vitro* condensates, and M. Turner, University of Warwick and H. Dale at the Molecular Imaging Centre, University of Bergen for insightful discussions. We also thank ARC at Durham University for the use of the Hamilton8 HPC service and EPCC at the University of Edinburgh for use of the Cirrus HPC service, which have both supported this work. S.N.G., H.K., T.A.W., T.S. and C.M.J. acknowledge funding from the Research Council of Norway (grant no. 335901). H.K. acknowledges funding from UKRI Engineering and Physical Sciences Research Council (grant no. EP/V034154/2). S.N.G. acknowledges support from the Trond Mohn Stiftelse (no. BFS2017TMT01) and The Royal Society, UK. S.N.G. and F.F.W acknowledge funding and support from the University of Bergen.

## 10 Author Contributions

J.O.L, T.A.W., C.M.J, F.W and E.S.T wrote the software. T.S. and T.A.W. performed the experiments. T.A.W. and F.W. performed the *FlickerPrint* analysis. H.K. and S.N.G supervised the project. T.A.W., J.O.L, T.S. and F.W. prepared the manuscript. H.K. and S.N.G. edited the manuscript. All authors were involved in discussions to develop the software, experiments and final manuscript.

## 12 Resource Availability

Source code for *FlickerPrint* is available on GitHub^45^. Documentation for *FlickerPrint* can be found at^40^. Code and raw data for producing the figures in this work will be released at the time of publication.

## References

1 Feric, M., Vaidya, N., Harmon, T. S., Mitrea, D. M., L. Zhu, Richardson, T. M., Kriwacki, R. W., Pappu, R. V., and Brangwynne, C. P. (2016). Coexisting liquid phases underlie nucleolar subcompartments. Cell 1, 1686–1697. doi:10.1016/j.cell.2016.04.047.

2 Banani, S. F., Lee, H. O., Hyman, A. A., and Rosen, M. K. (2017). Biomolecular condensates: organizers of cellular biochemistry. Nature Reviews Molecular Cell Biology 18, 285– 298. doi:10.1038/nrm.2017.7.

3 Kayali, F., Montie, H. L., Rafols, J. A., and Gracia, D. J. (2005). Prolonged translation arrest in reperfuesd hippocampal cornu ammonis 1 is mediated by stress granules. Cellular Neuroscience 134, 1223–1245. doi:10.1016/j.neuroscience.2005.05.047.

4 Correll, C. C., Bartek, J., and Dundr, M. (2019). The nucleolus: A multiphase condensate balancing ribosome synthesis and translational capacity in health, aging and ribosomopathies. Cells 8, 869. doi:10.3390/cells8080869.

5 Sharp, P. A., Chakraborty, A. K., Henninger, J. E., and Young, R. A. (2022). Rna in formation and regulation of transcriptional condensates. RNA (52–57). doi:10.1261/rna.078997.121.

6 Law, J. O., Jones, C. M., Stevenson, T., Williamson, T. A., Turner, M. S., Kusumaatmaja, H., and Grellscheid, S. N. (2023). A bending rigidity parameter for stress granule condensates. Science Advances 9, eadg0432. doi:10.1126/sciadv.adg0432.

7 Agudo-Canalejo, J., Schultz, S. W., Chino, H., Migliano, S. M., Saito, C., Koyama-Honda, I., Stenmark, H., Brech, A., May, A. I., Mizushima, N., and Knorr, R. L. (2021). Wetting regulates autophagy of phase-separated compartments and the cytosol. Nature 591, 142– 146. doi:10.1038/s41586-020-2992-3.

8 Lee, Y., Park, S., Yuan, F., Hayden, C. C., Wang, L., Lafer, E. M., Choi, S. Q., and Stachowiak, J. C. (2023). Transmembrane coupling of liquid-like protein condensates. Nature Communications 14, 8015. doi:10.1038/s41467-023-43332-w.

9 Galvanetto, N., Ivanović, M. T., Chowdhury, A., Sottini, A., Nü esch, M. F., Nettels, D., Best, R. B., and Schuler, B. (2023). Extreme dynamics in a biomolecular condensate. Nature 619, 876–883. doi:10.1038/s41586-023-06329-5.

10 Alshareedah, I., Borcherds, W. M., Cohen, S. R., Singh, A., Posey, A. E., Farag, M., Bremer, A., Strout, G. W., Tomares, D. T., Pappu, R. V., Mittag, T., and Banerjee, P. R. (2024). Sequence-specific interactions determine viscoelasticity and ageing dynamics of protein condensates. Nature Physics 20, 1482–1491. doi:10.1038/s41567-024-02558-1.

11 Alshareedah, I., Moosa, M. M., Pham, M., Potoyan, D. A., and Banerjee, P. R. (2021). Programmable viscoelasticity in protein-rna condensates with disordered sticker-spacer polypeptides. Nature Communications 12, 6620. doi:10.1038/s41467-021-26733-7.

12 Farag, M., Cohen, S. R., Borcherds, W. M., Bremer, A., Mittag, T., and Pappu, R. V. (2022). Condensates formed by prion-like low-complexity domains have small-world network structures and interfaces defined by expanded conformations. Nature Communications 13, 7722. doi:10.1038/s41467-022-35370-7.

13 Dai, Y., Chamberlayne, C. F., Messina, M. S., Chang, C. J., Zare, R. N., You, L., and Chilkoti, A. (2023). Interface of biomolecular condensates modulates redox reactions. Chem 9, 1594–1609. doi:10.1016/j.chempr.2023.04.001.

14 Molliex, A., Temirov, J., Lee, J., Coughlin, M., Kanagaraj, A. P., Kim, H. J., Mittag, T., and Taylor, J. P. (2015). Phase separation by low complexity domains promotes stress granule assembly and drives pathological fibrillization. Cell 163, 123–133. doi:10.1016/j.cell.2015.09.015.

15 Botterbusch, S., and Baumgart, T. (2021). Interactions between phase-separated liquids and membrane surfaces. Applied Sciences 11, 1288. doi:10.3390/app11031288.

16 Wiegand, T., and Hyman, A. A. (2020). Drops and fibers — how biomolecular condensates and cytoskeletal filaments influence each other. Emerging Topics in Life Sciences 4, 247– 261. doi:10.1042/ETLS20190174.

17 Shen, Y., Chen, A., Wang, W., Shen, Y., Ruggeri, F. S., Aime, S., Wang, Z., Qamar, S., Espinosa, J. R., Garaizar, A., George-Hyslop, P. S., Collepardo-Guevara, R., Weitz, D. A., Vigolo, D., and Knowles, T. P. J. (2023). The liquid-to-solid transition of fus is promoted by the condensate surface. Proceedings of the National Academy of Sciences 120, e2301366120. doi:10.1073/pnas.2301366120.

18 Choi, C.-H., Lee, D. S., Sanders, D. W., and Brangwynne, C. P. (2024). Condensate interfaces can accelerate protein aggregation. Biophysical Journal 123, 1404–1413. doi:10.1016/j.bpj.2023.10.009.

19 du Noüy, P.L. (1925). An interfacial tensiometer for universal use. Journal of General Physiology 7, 625–631. doi:10.1085/jgp.7.5.625.

20 Wilhelmy, L. (1863). Ueber die abhä ngigkeit der capillaritä ts-constanten des alkohols von substanz und gestalt des benetzten festen körpers. Annalen der Physik 195, 177–217. doi:10.1002/andp.18631950602.

21 Franses, E. I., Basaran, O. A., and Chang, C.-H. (1996). Techniques to measure dynamic surface tension. Current Opinion in Colloid & Interface Science 1, 296–303. doi:10.1016/S1359-0294(96)80018-5.

22 Caragine, C. M., Haley, S. C., and Zidovska, A. (2018). Surface fluctuations and coalescence of nucleolar droplets in the human cell nucleus. Phys. Rev. Lett. 121, 148101. doi:10.1103/PhysRevLett.121.148101.

23 Oranges, M., Jash, C., Golani, G., Seal, M., Cohen, S. R., Rosenhek-Goldian, I., Bogdanov, A., Safran, S., Daniella, and Goldfarb (2024). Core-shell model of the clusters of cpeb4 isoforms preceding liquid-liquid phase separation. Biophysical Journal 123, 2604–2622. doi:10.1016/j.bpj.2024.06.027.

24 Abbas, M., Law, J. O., Grellscheid, S. N., Huck, W. T. S., and Spruijt, E. (2022). Peptide-based coacervate-core vesicles with semipermeable membranes. Advanced Materials 34, 2202913. doi:10.1002/adma.202202913.

25 Helfrich, W. (1973). Elastic properties of lipid bilayers: Theory and possible experiments. Zeitschrift fü r Naturforschung C 28, 693–703. doi:DOI:10.1515/znc-1973-11-1209.

26 Milner, S. T., and Safran, S. A. (1987). Dynamical fluctuations of droplet microemulsions and vesicles. Physical Review A 36. doi:10.1103/PhysRevA.36.4371.

27 Brochard, F., and Lennon, J. (1975). Frequency spectrum of the flicker phenomenon in erythrocytes. Journal de Physique 36, 1035–1047. doi:10.1051/jphys:0197500360110103500.

28 Servuss, R. M., Harbich, V., and Helfrich, W. (1976). Measurement of the curvature-elastic modulus of egg lecithin bilayers. Biochimica et Biophysica Acta (BBA) - Biomembranes 436, 900–903. doi:10.1016/0005-2736(76)90422-3.

29 Fricke, K., Wirthensohn, K., Laxhuber, R., and Sackmann, E. (1986). Flicker spectroscopy of erythrocytes. European Biophysics Journal (67–81). doi:10.1007/BF00263063.

30 Yoon, Y.-Z., Hong, H., Brown, A., Kim, D. C., Kang, D. J., Lew, V. L., and Cicuta, P. (2009). Flickering analysis of erythrocyte mechanical properties: Dependence on oxygenation level, cell shape, and hydration level. Biophysical Journal 97, 1606–1615. doi:10.1016/j.bpj.2009.06.028.

31 Marr, D., and Hildreth, E. (1980). Theory of edge detection. Proceedings of the Royal Society of London. Series B. Biological Sciences 207, 187–217. doi:10.1098/rspb.1980.0020.

32 Pécréaux, J., Döbereiner, H.-G., Prost, J., Joanny, J.-F., and Bassereau, P. (2004). Refined contour analysis of giant unilamellar vesicles. The European Physical Journal E 13, 277– 290. doi:10.1140/epje/i2004-10001-9.

33 Schindelin, J., Arganda-Carreras, I., Frise, E., Kaynig, V., Longair, M., Pietzsch, T., Preibisch, S., Rueden, C., Saalfeld, S., Schmid, B., Tinevez, J.-Y., White, D. J., Hartenstein, V., Eliceiri, K., Tomancak, P., and Cardona, A. (2012). Fiji: an open-source platform for biological-image analysis. Nature Methods 9, 676=682. doi:10.1038/nmeth.2019.

34 Arter, W. E., Qi, R., Erkamp, N. A., Krainer, G., Didi, K., Welsh, T. J., Acker, J., Nixon-Abell, J., Qamar, S., Guillén-Boixet, J., Franzmann, T. M., Kuster, D., Hyman, A. A., Borodavka, A., George-Hyslop, P. S., Alberti, S., and Knowles, T. P. J. (2022). Biomolecular condensate phase diagrams with a combinatorial microdroplet platform. Nature Communications 13, 7845. doi:10.1038/s41467-022-35265-7.

35 Dörner, K., Gut, M., Overwijn, D., Cao, F., Siketanc, M., Heinrich, S., Beuret, N., Sharpe, T., Lindorff-Larsen, K., and Maria, H. (). Tag with caution — how protein tagging influences the formation of condensates. bioRxiv. doi:10.1101/2024.10.04.616694.

36 Graham, K., Chandrasekaran, A., Wang, L., Ladak, A., Lafer, E. M., Rangamani, P., and Stachowiak, J. C. (2023). Liquid-like vasp condensates drive actin polymerization and dynamic bundling. Nature Physics 19, 574–585. doi:10.1038/s41567-022-01924-1.

37 Walker, C., Chandrasekaran, A., Mansour, D., Graham, K., Torres, A., Wang, L., Lafer, E. M., Rangamani, P., and Stachowiak, J. C. (2025). Liquid-like condensates that bind actin promote assembly and bundling of actin filaments. Developmental Cell 60, 1–18. doi:10.1016/j.devcel.2025.01.012.

38 Döbereiner, H.-G. (2000). Properties of giant vesicles. Current Opinion in Colloid & Interface Science 5, 256–263. doi:10.1016/S1359-0294(00)00064-9.

39 Rautu, S. A., Orsi, D., Di Michele, L., Rowlands, G., Cicuta, P., and Turner, M. S. (2017). The role of optical projection in the analysis of membrane fluctuations. Soft Matter 13, 3480–3483. doi:10.1039/C7SM00108H.

40 FlickerPrint (2025). Flickerprint documentation. https://flickerprint.github.io/FlickerPrint/.

41 Wilkinson, M. D., Dumontier, M., Aalbersberg, I. J., Appleton, G., Axton, M., Baak, A., Blomberg, N., Boiten, J.-W., da Silva Santos, L. B., Bourne, P. E., Bouwman, J., Brookes, J., Clark, T., Crosas, M., Dillo, I., Dumon, O., Edmunds, S., Evelo, C. T., Finkers, R., Gonzalez-Beltran, A., Gray, A. J., Groth, P., Goble, C., Grethe, J. S., Heringa, J., ‘t Hoen, P. A., Hooft, R., Kuhn, T., Kok, R., Kok, J., Lusher, S. J., Martone, M. E., Mons, A., Packer, A. L., Persson, B., Rocca-Serra, P., Roos, M., van Schaik, R., Sansone, S.-A., Schultes, E., Sengstag, T., Slater, T., Strawn, G., Swertz, M. A., Thompson, M., van der Lei, J., van Mulligen, E., Velterop, J., Waagmeester, A., Wittenburg, P., Wolstencroft, K., Zhao, J., and Mons, B. (2016). The fair guiding principles for scientific data management and stewardship. Scientific Data 3, 160018. doi:10.1038/sdata.2016.18.

42 Linkert, M., Rueden, C. T., Allan, C., Burel, J.-M., Moore, W., Patterson, A., Loranger, B., Moore, J., Neves, C., MacDonald, D., Tarkowska, A., Sticco, C., Hill, E., Rossner, M., Eliceiri, K. W., and Swedlow, J. R. (2010). Metadata matters: access to image data in the real world. Journal of Cell Biology 189, 777–782.

43 Picheny, V., Berkeley, J., Moss, H. B., Stojic, H., Granta, U., Ober, S. W., Artemev, A., Ghani, K., Goodall, A., Paleyes, A., Vakili, S., Pascual-Diaz, S., Markou, S., Qing, J., Loka, N. R. B. S., Couckuyt, I., and Morter, C. (2023). Trieste: Efficiently exploring the depths of black-box functions with tensorflow. arXiv. URL: https://arxiv.org/abs/2302.08436. doi:10.48550/ARXIV.2302.08436.

44 Gardner, J. R., Kusner, M. J., Xu, Z. E., Weinberger, K. Q., and Cunningham, J. P. (2014). Bayesian optimization with inequality constraints.

45 FlickerPrint (2025). Flickerprint github repository. https://github.com/FlickerPrint/ FlickerPrint.

46 The HDF Group (). Hierarchical data format, version 5. https://www.hdfgroup.org/HDF5/.

47 Fabrini, G., Farag, N., Nuccio, S. P., Li, S., Stewart, J. M., Tang, A. A., McCoy, R., Owens, R. M., Rothemund, P. W. K., Franco, E., Di Antonio, M., and Di Michele, L. (2024). Cotranscriptional production of programmable rna condensates and synthetic organelles. Nature Nanotechnology 19, 1665–1673. doi:10.1038/s41565-024-01726-x.

48 Kedersha, N., Panas, M. D., Achorn, C. A., Lyons, S., Tisdale, S., Hickman, T., Thomas, M., Lieberman, J., McInerney, G. M., Ivanov, P., and Anderson, P. (2016). G3bp–caprin1–usp10 complexes mediate stress granule condensation and associate with 40s subunits. Journal of Cell Biology 212, e201508028. doi:10.1083/jcb.201508028.

49 Yewdall, N. A., André, A. A., van Haren, M. H., Nelissen, F. H., Jonker, A., and Spruijt, E. (2022). Atp:mg^2+^ shapes material properties of protein-rna condensates and their partitioning of clients. Biophysical Journal 121. doi:10.1016/j.bpj.2022.08.025.

