## Supplemental Material for "FlickerPrint: An Analysis Package for Measuring Interfacial Tension and Bending Rigidity of Biomolecular Condensates and Vesicles at Scale"

### SM1: Object Tracking

As discussed in the main text, in each frame of the input microscope file, objects of interest (condensates or vesicles) are detected using the Difference of Gaussians method[1]. The next step is to match objects between frames, so that per-object averages can be calculated at later steps. This tracking process poses some unique challenges in the case of *FlickerPrint*. Here, we outline the bespoke tracking algorithm used by *FlickerPrint*.

The greatest challenge for designing the tracking algorithm is that the Difference of Gaussians detector is not fully reliable, which means that an object may be missed for a few frames, from time to time. Therefore judgments must be made about whether a newly detected object is genuinely newly-appeared, or whether it is a previously detected object that has been dropped for a few frames. As a result, the tracking does not only match to objects that occurred on the previous frame, but retains a memory of the last known position of each object, that endures for 10 frames.

Conversely, as the objects should all lie on a plane, and we do not expect them to move very quickly (and ideally not at all), we are absolved from the need to deal with crossing trajectories or ballistic motion. Therefore, we have designed a tracking algorithm that is based on object positions only; ignoring velocity or shape-matching.

The *FlickerPrint* tracking algorithm is described below:

1. On the first frame, there is no need for tracking. Each object detected is assigned a unique integer ID, starting from 0 and counting up. The position and ID of each object is stored in the tracker's memory for comparison with future frames.

2. On subsequent frames, we begin by looking for objects that appear both in the tracker's memory and the current frame. An object in the current frame is identified as an object in the tracker's memory if their centres are within 15 pixels of each other, and they are closer than any other object. The ID of the object in the current frame is set to the ID stored in the tracker's memory, and the position stored in the tracker's memory is updated to the latest position. If the object is one that was previously missing, then its countdown is stopped and deleted (see step 4.).

3. This leaves two classes of unhandled object: objects that appear in the current frame but not in the tracker's memory, and objects that appear in the tracker's memory but not the current frame. The first class are newly appeared objects. They are assigned a currently-unused ID and added in to the tracker's memory.

4. The other class are the missing objects. If an object is newly missing (i.e. it appeared in the previous frame), a countdown is started from 10, but the object remains in the tracker's memory at its last known position. If the object already has a countdown (i.e. it has been missing for more than one frame), then the countdown is decremented by one. If the countdown reaches zero the object is deleted from the memory permanently. Therefore, if an object subsequently appears at that position, it will be counted as a new object.

The memory value of 10 frames is chosen so that an object that is missed by the detector for a few frames is usually re-identified, but that it is very rare that a granule will drift onto the position of a previously-vanished object and be miss-identified as the same object.

#### SM2: Boundary Detection

In order to estimate the boundary of the objects of interest, we must first determine the extent of the object in the frame. Using the centre point found by the difference of Gaussians algorithm, we use a simple flood fill to estimate the extent of the object, and use this as a guide to cut out a rectangular region containing just that object. The extent of the flood fill can be set in the configuration file.

We then smooth the image by applying a Gaussian blur with a width set in the configuration file. This reduces internal gradients caused my noise in the microscope file. The procedures described above are the same for both vesicles and condensates. However, at this point condensates require an additional processing step. We take the directional gradient $g = \nabla I \cdot \hat{\mathbf{r}}$, where $I$ is the image and $\hat{\mathbf{r}}$ is the radial unit vector field from the condensate centre. We estimate $\nabla I$ using a fourth order kernel, given by $\mathbf{k}_{4\text{th}} = \frac{1}{12}\begin{pmatrix} 1 & -8 & 0 & 8 & -1 \end{pmatrix}$. This kernel is applied to the input image as a stencil, centred at each pixel, and multiplied element-wise[2]. As we expect $g$ to be highest at the edge of the condensate, the following steps are once again the same for vesicles and condensates.

To find the outline of the object, we emit 400 rays radially from the centre point of the object. We sample the processed image intensity at equally spaced points along these rays at a density 15 per pixel-width. We use a fourth-order interpolation to obtain values resolved at a sub-pixel level. For each ray, we take the sampled point with the highest intensity value. This leaves with 400 points on the frame that describe the outline of the object.

#### SM3: Justification of the resolution of boundary detection

As described in the main text and in SM2, for objects with appear as solid regions of high intensity (rather than high intensity outlines) such as condensates (main text Figure 2 e,f), the boundary is taken as the contour of maximum intensity gradient, measured radially from the centre of the object. The condensate boundary can be determined to a resolution of $\frac{1}{15}$ of a pixel. Here, we justify this claim.

The normalised 1-Dimensional intensity profile of the boundary of a condensate can be represented by

$$I(x) = 1/2 - 1/2 \tanh\left(\frac{x - x_0}{\zeta}\right) \tag{S1}$$

where $x$ is an integer representing the position of a pixel, $x_0$ is the 'true' position of the boundary and $\zeta$ is the interface width; typically $\zeta = 0.8$. An image with 8-bit colour-depth contains 255 intensity levels per pixel. Requiring a difference of 10 levels in order to determine the boundary position ensures that the intensity gradient is sufficiently steep to avoid mis-detections and provides allowances for when the contrast in an image is not optimal. Therefore, the difference in the intensity of a pixel at $x$ when the boundary is at locations $x$ and $x + \alpha$ is

$$\frac{1}{2}\tanh\left(\frac{x - x}{\zeta}\right) - \frac{1}{2}\tanh\left(\frac{x - (x + \alpha)}{\zeta}\right) = \frac{n_{levels}}{255}. \tag{S2}$$

Simplifying down gives

$$\tanh\left(\frac{\alpha}{\zeta}\right) = 2\frac{n_{levels}}{255}. \tag{S3}$$

Substituting in values gives $\alpha = \frac{1}{15.9}$. Thus, we determine the condensate boundary to a resolution of 1/15 of a pixel.

### SM4: Spectrum Fitting

The final step in calculating the interfacial tension and bending rigidity of a condensate or vesicle is fitting experimentally measured fluctuation spectrum (main text, figure 1 F) to Equation 1 of the main text. Firstly, we must define an error function $\varepsilon$, which allows us to compare the quality of fits. We previously showed that the best error function for this problem is

$$\varepsilon = \sum_q \left| \log_{10} \left[ \frac{|F_{q,\mathrm{theo}}|^2}{|F_{q,\mathrm{exp}}|^2} \right] \right|, \tag{S4}$$

where $|F_{q,\mathrm{theo}}|$ is the theoretical fluctuation spectrum and $|F_{q,\mathrm{exp}}|$ is the experimental spectrum[3]. This value is minimised to find the best fit for $\sigma$ and $\kappa$.

Figure S3 shows a representative fitting surface for a stress granule, across several orders of magnitude. It can be seen that there is only one minimum, which means that down-hill fitting methods are sufficient to find the global minimum. Therefore, fitting is carried out as follows:

1. We generate an initial guess by sampling $\bar{\sigma}$ and $\kappa$ across 14 orders of magnitude on a logarithmic scale. The pair that gives the lowest $\varepsilon$ is used as a starting point for the next step.
2. We then use the least_squares function from the scipy.optimize python module to find the true minimum of $\varepsilon$ [4].

### SM5: Robustness of *FlickerPrint* Analysis to Changes in Imaging Parameters

As discussed in the main text, *FlickerPrint* contains five parameters which can be adjusted to account for differences in imaging setup. These are the minimum intensity (relative to the frame maximum intensity) which an object must have in order to be detected, its approximate maximum and minimum size, the threshold intensity of the flood fill and a smoothing parameter which is used to account for microscope noise. Choosing appropriate parameters ensures that the maximum number of objects of interest are found in each frame, whilst minimising the number of false positive detections. However, here we show that the results of *FlickerPrint* analysis are relatively robust to changes in these parameters.

Figure S4 shows the effect of systematically varying each of the imaging parameters for an image of stress granules in U2OS cells. . In each case, the final number of condensates analysed after the parameter distributions have been filtered (Methods, main paper), and the returned distributions of interfacial tension, bending rigidity and mean condensate radius are plotted. When allowing the imaging parameters to be varied across a wide range of values, interfacial tension, bending rigidity and mean radius vary by up to 1.3 $N/m$, 12 $k_BT$ and 0.75 $\mu m$ respectively. To allow for direct comparison, it can be helpful to quote the variation in the population distributions in terms of their geometric standard deviation (for interfacial tension and bending rigidity) and standard deviation (for mean radius). When quoted as a fraction of the standard deviation of the two parameter distributions in least agreement, interfacial tension, bending rigidity and mean radius vary by $< 1.3$ SD, $< 4.1$ SD and $< 15.3$ SD respectively (Figure S4). However, when the parameters are within their optimal range (defined as the approximate range of parameters which maximise the number of detected objects, whilst minimising the number of objects which fail the later filtering steps; see SM6), the variation in the interfacial tension, bending rigidity and mean radius distributions for all condensates is reduced to $<$ 0.52 SD, $<$ 0.42 SD and $<$ 0.93 SD respectively.

Comparing the final population distributions can be useful to gauge the variation in population distributions caused by changes in the imaging parameters. However, since changing the imaging parameters affects the number of condensates which are found, comparing the populations of all condensates does not provide much information on how the properties of individual condensates are affected. Therefore, Figure S4 A-D also shows the returned parameter distributions when only condensates which are found and pass the filters in at least 75% of analyses are included. A cut-off of 75% ensures that the distributions contain broadly the same condensates, whilst allowing flexibility to ensure that the population distributions are large enough for statistical analysis. In this case, the parameter distributions for interfacial tension, bending rigidity and mean radius deviate by $<$ 0.89SD, $<$ 1.1SD and $<$ 1.9SD respectively across the full range of imaging parameters, and $<$ 0.28SD, $<$ 0.45SD and $<$ 0.12SD respectively when only optimal parameters are considered. These variations in the returned distributions are much less than for the case where all condensates are considered. This supports the idea that the variations are largely due to different condensates being found when the imaging parameters are changed, since these parameters control where objects are found in the image, not the position of their boundary.

Once objects have been found, their boundary is determined as described in SM2. This process is configured by a single parameter which smooths the image to account for microscope noise which would cause discontinuities in the object boundary. Similar analysis to that conducted for the parameters responsible for object location has been conducted for the smoothing parameter and is shown in Figure S4 E. In this case, the typical variation in the returned parameter distributions across the full range of smoothing values is small compared with the imaging parameters used for locating objects ($< 0.61$ SD, $< 0.31$ SD and $< 2.3$ SD for interfacial tension, bending rigidity and mean radius). However, the distributions from condensates found in at least 75% of analyses do not see the same reduction in variation as for the other imaging parameters ($< 0.41$ SD, $< 0.76$ SD and $< 0.87$ SD for interfacial tension, bending rigidity and mean radius). This difference in behaviour is likely a result of the shape of the condensate boundary being directly impacted by the value of the smoothing parameter. However, as discussed in the main text, the smoothing parameter should typically be kept close to 1.0 and should only be increased when microscope noise cannot be reduced using other methods.

### SM6: Bayesian Optimisation Tool for Image Parameter Estimation

Although we have shown in SM5 that the analysis performed by *FlickerPrint* is robust to changes in imaging parameters, it is still important to ensure that optimal parameters are selected. A good set of imaging parameters result in a high number of valid objects of interest (objects which pass all filtering steps) and a low number of rejected objects (those which do not pass one of the filters for a given frame). To help with choice of parameters, *FlickerPrint* contains an optimisation tool to determine the optimal imaging parameters (minimum object intensity, and maximum and minimum object radius) to maximise the number of condensates found in an assay.

In order to do so, it is necessary to quantify how good a given set of parameters is. For this, two objectives are used: i) the number of usable condensates found need to be as high as possible and ii) the number of unusable condensates should be as low as possible. Typically, these two objectives do not align with each other; if the number of usable condensates is high, then the number of unusable condensates is often also high. Conversely, if the number of unusable condensates is low, the number of usable condensates is typically also low. It is therefore necessary to balance these two objectives against each other.

As a given assay often contains multiple videos, each with hundreds of frames, repetitive evaluation of every frame in every video would be very computationally expensive. Therefore, only a sample set of frames are used for the parameter estimation. The selected frames are taken from a subset, $E_{\text{sub}}$, of up to 12 videos, sampled from a uniform distribution of all the provided videos. For each video 2 random frames, $I_{\text{sub}}$ are analysed, again sampled from a uniform distribution.

If $N$ is the number of videos in $E_{\text{sub}}$, $M$ is the number of frames in $I_{\text{sub}}$ and $\theta$ is a set of imaging parameters parameters, then the two objectives can be written as

$$\mathcal{F}_1(\theta, E_{\text{sub}}) = \sum_{I_{\text{sub}} \in E_{\text{sub}}} \min_{f \in I_{\text{sub}}} \mathrm{DoG}_{\text{usable}}(\theta, f), \tag{S5}$$

and

$$\mathcal{F}_2(\theta, E_{\text{sub}}) = -\frac{1}{N} \sum_{I_{\text{sub}} \in E_{\text{sub}}} \left( \frac{1}{M} \sum_{f \in I_{\text{sub}}} \mathrm{DoG}_{\text{unusable}}(\theta, f) \right), \tag{S6}$$

where $\mathrm{DoG}_{\text{usable}}$ and $\mathrm{DoG}_{\text{unusable}}$ are instances of the Difference of Gaussians algorithm that return the number of usable and unusable objects in frame $f$, respectively.

$\mathcal{F}_1$ corresponds to objective i) and tries to maximize the sum of the number of usable objects of interest for the frame with the fewest number of usable objects in each image. This approach is chosen to find a parameter set where the floor of found condensates is increased as much as possible. This has the benefit that objects are present in as many images as possible, allowing for better analysis with *FlickerPrint*.

$\mathcal{F}_2$ corresponds to objective ii) and tries to minimize the mean number of unusable objects of interest over all images, where each image is weighted by the mean number of unusable objects found in its frames. Contrary to objective i), it is not necessary to worry about continuous existence of the objects, only the total number. The goal is therefore to try and find a parameter set that reduces the mean number of unusable objects of interest.

Since *FlickerPrint* performs best with a high number of usable condensates, objective i) is prioritized over objective ii). Using this, the problem can be described as a multi-objective optimisation problem with ranked objectives[5]. The problem can be written as a lexicographic optimisation

$$
\begin{aligned}
\operatorname{lex\,max} \quad & \mathcal{F}_1(\theta; E_{\text{sub}})\mathcal{F}_2(\theta, E_{\text{sub}}) && \text{(S7)}\\
\text{s.t.} \quad & \theta \in \Theta, && \text{(S7a)}
\end{aligned}
$$

where $\Theta$ is the set of all possible imaging parameters. EquationS7 is solved with a sequential algorithm, meaning that the problem is divided into two single-objective optimisation problems which are solved sequentially.

The first problem is given by

$$
\begin{aligned}
n_{\text{max}} = \max \quad & \mathcal{F}_1(\theta; E_{\text{sub}}) && \text{(S8)}\\
\text{s.t.} \quad & \theta \in \Theta && \text{(S8a)}
\end{aligned}
$$

with $n_{max}$ being the maximum number of usable condensates found in $E_{\text{sub}}$.
The second optimisation problem is given by

$$
\begin{aligned}
\max \quad & \mathcal{F}_2(\theta; E_{\text{sub}}) && \text{(S9)}\\
\text{s.t.} \quad & \theta \in \Theta, && \text{(S9a)}\\
& \mathcal{F}_1(\theta; E_{\text{sub}}) \geq 0.9 n_{\text{max}}. && \text{(S9b)}
\end{aligned}
$$

The second optimisation is constrained by the maximum number of usable condensates found in the first, higher prioritized optimization. While the ideal choice of the constraint is dependent on the assay, we found that $0.9$ resulted in good results in most tested cases.
Since the evaluation of $\mathcal{F}_1$ and $\mathcal{F}_2$ is of high cost and since $\text{DoG}_{\text{usable}}$ and $\text{DoG}_{\text{unusable}}$ do not take on a functional form, Bayesian optimization is used to solve equation S7[6,7].
Figure S5 demonstrates an application of the Bayesian optimisation tool to stress granules induced in U2OS cells; Figure S5 A shows that the optimisation tool is able to successfully converge on a set of parameters $\theta$ to minimise the objective function. The variation seen in the start and end points of the optimisation are due to different frames being selected for each repeat. However, as demonstrated in Figure S5 B, these differences lead to minimal variation in the optimised parameters. Taken together with the analysis conducted in SM5, the optimisation tool yields stable distributions of the mechanical properties of the objects of interest at the population level.

### Supplementary Figures

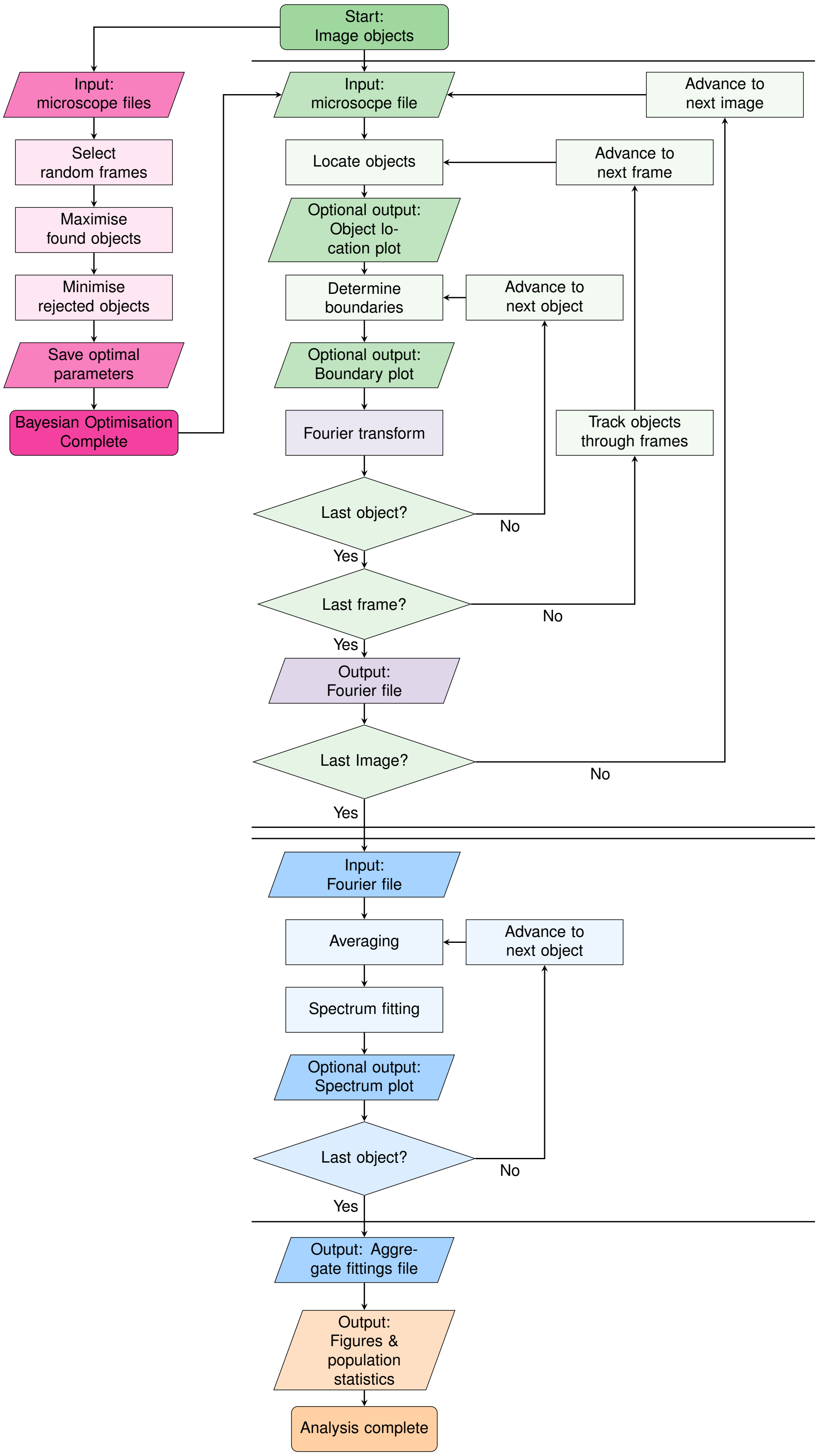


Figure S1: **A flowchart detailing the full workflow of *FlickerPrint*.** The workflow is split into five stages; location of granules in a microscope image (green), Fourier Transform of the boundary fluctuations (purple), fitting of the theoretical power spectrum (blue), Bayesian parameter estimation (pink) and analysis of population-level statistics (orange). Steps in between parallel lines indicate that work on multiple microscope images can be parallelised across multiple cores.

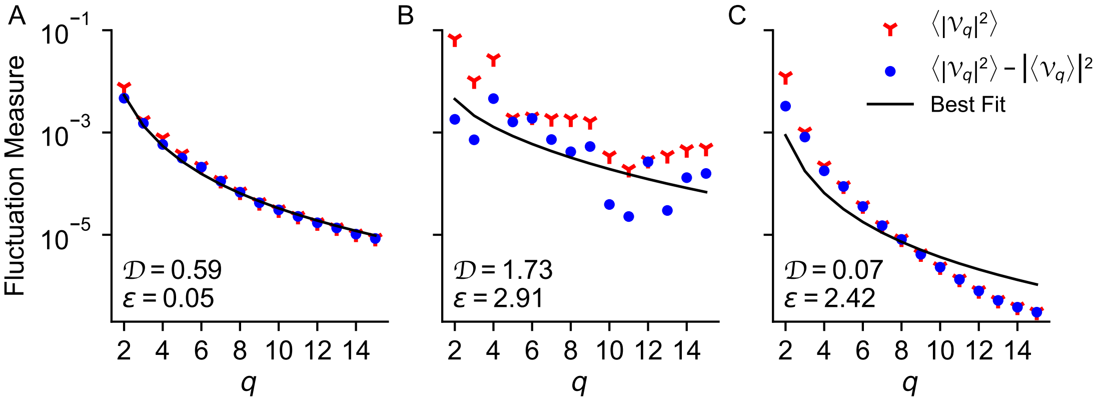


Figure S2: **The fitting error $\varepsilon$ and Durbin-Watson statistic $\mathcal{D}$ can be used as a measure of the goodness of fit of the analytic power spectrum to the measured fluctuation amplitudes[8].** Three example spectra of stress granules in U2OS cells are shown. In (**A**), the model is a good fit to the experimental spectrum; in (**B**), the experimental spectrum is very noisy and in (**C**), the model does not fit the experimental spectrum well.

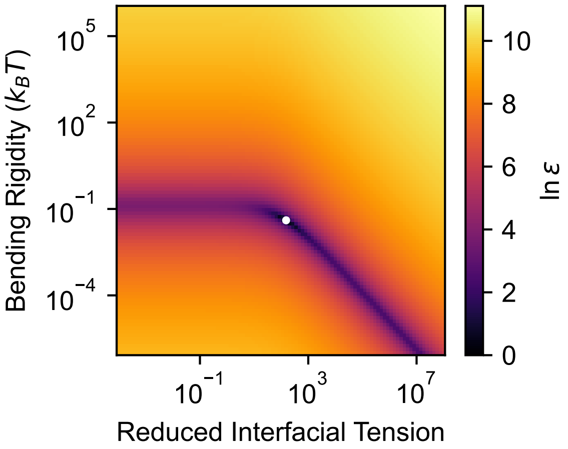


Figure S3: **A heat-map showing the error surface $\varepsilon(\bar{\sigma}, \kappa)$ for a representative stress granule.** The minimum of the surface is indicated with a white dot. The $x$-axis shows reduced interfacial tension $\bar{\sigma} = \frac{\sigma R^2}{\kappa}$.

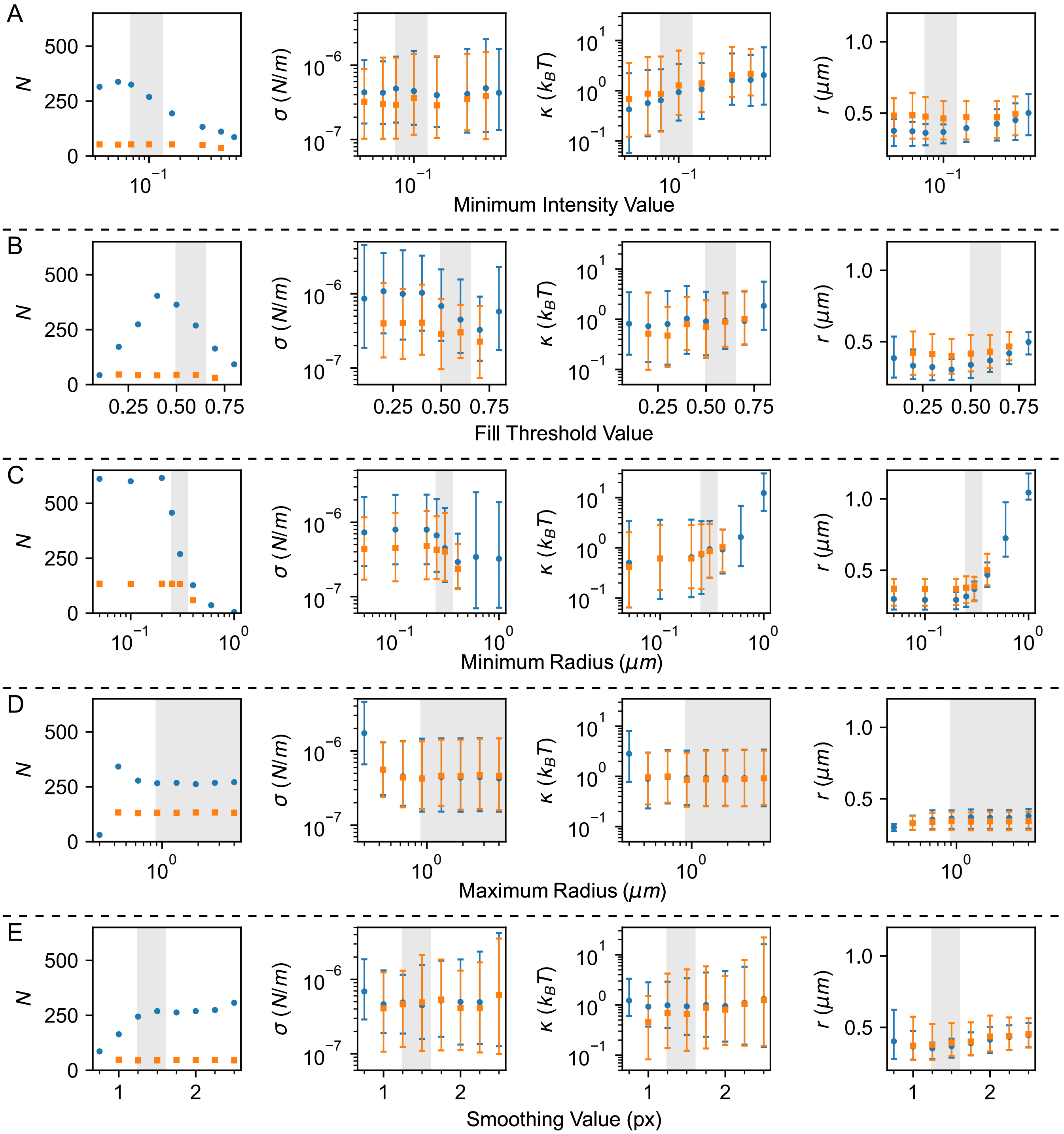


Figure S4: **The parameter distributions output by *FlickerPrint* are robust to variations in the imaging parameters used to configure the analysis.** The values for minimum object intensity (**A**), flood fill threshold (**B**), minimum object radius (**C**), maximum radius (**D**) and the smoothing parameter (**E**) are varied independently for analysis of one video of stress granules in U2OS cells. The effect on the number of condensates which are found and pass the filtering stages $N$, the interfacial tension $\sigma$, bending rigidity $\kappa$ and mean condensate radius $r$ are plotted. Points show the mean value of the output distributions; error bars show $\pm$ 1 standard deviation. Blue circles show data for all condensates which pass the filtering steps; orange squares show distributions which only contain condensates which are consistently found across 75% of analyses of the same parameter. Where less than 100 condensates from one dataset passed the filters, tracking between datasets was not performed. Light grey regions indicate the window of optimal parameters for the image analysed.

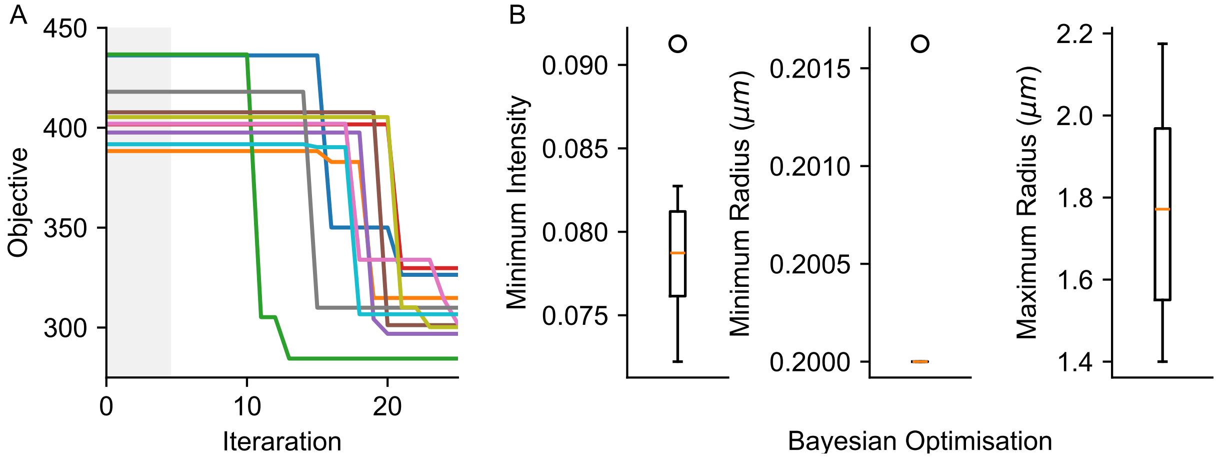


Figure S5: **Bayesian Optimisation can be used to determine appropriate imaging parameters for *FlickerPrint* analysis.** (**A**) Value of the objective function given by equation S7 with iteration count for the lexicographic optimisation, performed on images of stress granules in U2OS cells. 10 repeats, each using 5 frames from 3 randomly selected images are shown. The shaded grey region is the 'burn in' period (5 iterations) for the optimisation. (**B**) Box plots showing the spread of optimised imaging parameters (minimum object intensity, minimum radius and maximum radius), as determined by the optimisations performed in (A).

### Supplementary Tables

| Frame Rate (fps) | Number of Condensates | Interfacial Tension $^{+1SD}_{-1SD}$ $(\mu N/m)$ | Bending Rigidity $^{+1SD}_{-1SD}$ $(k_BT)$ | Mean Radius $^{+1SD}_{-1SD}$ $(\mu m)$ |
|---|---|---|---|---|
| 6 | 121 | $0.259^{+0.326}_{-0.135}$ | $0.92^{+1.44}_{-0.53}$ | $0.40^{+0.08}_{-0.09}$ |
| 8 | 121 | $0.281^{+0.394}_{-0.161}$ | $0.92^{+1.59}_{-0.56}$ | $0.40^{+0.08}_{-0.10}$ |
| 12 | 121 | $0.297^{+0.331}_{-0.160}$ | $0.94^{+1.32}_{-0.55}$ | $0.40^{+0.08}_{-0.10}$ |
| 24 | 121 | $0.302^{+0.428}_{-0.171}$ | $0.91^{+1.38}_{-0.51}$ | $0.40^{+0.08}_{-0.10}$ |

Table S1: **Statistics for parameter distributions of the same two videos of stress granules in U2OS cells, analysed at differing effective frame rates.** Interfacial tension and bending rigidity distributions are shown in Figure 5 A, B of the main text. Only condensates which are found, analysed and pass the filtering steps are included in the statistics. Consequently, there is very little deviation in the parameter distributions.

| Frame Rate (fps) | Number of Condensates | Interfacial Tension $^{+1SD}_{-1SD}$ $(\mu N/m)$ | Bending Rigidity $^{+1SD}_{-1SD}$ $(k_BT)$ | Mean Radius $^{+1SD}_{-1SD}$ $(\mu m)$ |
|---|---|---|---|---|
| 6 | 177 | $0.299^{+0.581}_{-0.173}$ | $0.93^{+1.83}_{-0.0.63}$ | $0.38^{+0.05}_{-0.08}$ |
| 8 | 227 | $0.316^{+0.442}_{-0.179}$ | $0.88^{+1.76}_{-0.56}$ | $0.36^{+0.04}_{-0.07}$ |
| 12 | 268 | $0.367^{+0.679}_{-0.235}$ | $1.06^{+2.37}_{-0.72}$ | $0.36^{+0.05}_{-0.07}$ |
| 24 | 400 | $0.417^{+0.646}_{-0.253}$ | $1.07^{+2.07}_{-0.72}$ | $0.34^{+0.03}_{-0.06}$ |

Table S2: Similar to Table S1, but statistics are for all condensates found in each set of analysis, regardless of whether they were found in other analyses. More condensates can be tracked successfully and are visible in sufficiently many frames for their fluctuations to be averaged at higher frame rates. This leads to variations in the parameter distributions.

### References


1. Marr, D., and Hildreth, E. (1980). Theory of edge detection. Proceedings of the Royal Society of London. Series B. Biological Sciences *207*, 187–217. doi:10.1098/rspb.1980.0020.

2. Jones, C. Developing computational tools to investigate stress granules *In-vivo*. Ph.D. thesis Durham University (2020).

3. Law, J. O., Jones, C. M., Stevenson, T., Williamson, T. A., Turner, M. S., Kusumaatmaja, H., and Grellscheid, S. N. (2023). A bending rigidity parameter for stress granule condensates. Science Advances *9*, eadg0432. doi:10.1126/sciadv.adg0432.

4. Virtanen, P., Gommers, R., Oliphant, T. E., Haberland, M., Reddy, T., Cournapeau, D., Burovski, E., Peterson, P., Weckesser, W., Bright, J., van der Walt, S. J., Brett, M., Wilson, J., Millman, K. J., Mayorov, N., Nelson, A. R. J., Jones, E., Kern, R., Larson, E., Carey, C. J., Polat, İ., Feng, Y., Moore, E. W., VanderPlas, J., Laxalde, D., Perktold, J., Cimrman, R., Henriksen, I., Quintero, E. A., Harris, C. R., Archibald, A. M., Ribeiro, A. H., Pedregosa, F., van Mulbregt, P., and SciPy 1.0 Contributors (2020). SciPy 1.0: Fundamental Algorithms for Scientific Computing in Python. Nature Methods *17*, 261–272. doi:10.1038/s41592-019-0686-2.

5. Marler, R., and Arora, J. (2004). Survey of multi-objective optimization methods for engineering. STRUCTURAL AND MULTIDISCIPLINARY OPTIMIZATION *26*, 369–395. doi:10.1007/s00158-003-0368-6.

6. Picheny, V., Berkeley, J., Moss, H. B., Stojic, H., Granta, U., Ober, S. W., Artemev, A., Ghani, K., Goodall, A., Paleyes, A., Vakili, S., Pascual-Diaz, S., Markou, S., Qing, J., Loka, N. R. B. S., Couckuyt, I., and Morter, C. (2023). Trieste: Efficiently exploring the depths of black-box functions with tensorflow. arXiv. URL: https://arxiv.org/abs/2302.08436. doi:10.48550/ARXIV.2302.08436.

7. Gardner, J. R., Kusner, M. J., Xu, Z. E., Weinberger, K. Q., and Cunningham, J. P. (2014). Bayesian optimization with inequality constraints.

8. Durbin, J., and Watson, G. S. (1950). Testing for serial correlation in least squares regression: I. Biometrika *37*, 409–428. doi:10.2307/2332391.
